# Inferring sparse structure in genotype-phenotype maps

**DOI:** 10.1101/2022.09.27.509675

**Authors:** Samantha Petti, Gautam Reddy, Michael M. Desai

**Affiliations:** NSF-Simons Center for the Mathematical and Statistical Analysis of Biology, Harvard University, Cambridge, MA, USA; NSF-Simons Center for the Mathematical and Statistical Analysis of Biology, Harvard University. Physics & Informatics Laboratories, NTT Research, Inc., Sunnyvale, CA, USA; Center for Brain Science, Harvard University, Cambridge, MA, USA; Organismic and Evolutionary Biology and Physics, Harvard University, Cambridge, MA, USA

## Abstract

Correlation among multiple phenotypes across related individuals may reflect some pattern of shared genetic architecture: individual genetic loci affect multiple phenotypes (an effect known as pleiotropy), creating observable relationships between phenotypes. A natural hypothesis is that pleiotropic effects reflect a relatively small set of common “core” cellular processes: each genetic locus affects one or a few core processes, and these core processes in turn determine the observed phenotypes. Here, we propose a method to infer such structure in genotype-phenotype data. Our approach, *sparse structure discovery* (SSD) is based on a penalized matrix decomposition designed to identify latent structure that is low-dimensional (many fewer core processes than phenotypes and genetic loci), locus-sparse (each locus affects few core processes) and/or phenotype-sparse (each phenotype is influenced by few core processes). Our use of sparsity as a guide in the matrix decomposition is motivated by the results of a novel empirical test indicating evidence of sparse structure in several recent genotype-phenotype data sets. First, we use synthetic data to show that our SSD approach can accurately recover core processes if each genetic locus affects few core processes or if each phenotype is affected by few core processes. Next, we apply the method to three datasets spanning adaptive mutations in yeast, genotoxin robustness assay in human cell lines, and genetic loci identified from a yeast cross, and evaluate the biological plausibility of the core process identified. More generally, we propose sparsity as a guiding prior for resolving latent structure in empirical genotype-phenotype maps.

## I. INTRODUCTION

A central goal of quantitative genetics is to exploit observed correlations between genotype and phenotype to infer the structure of the genotype-phenotype map [1–6]. That is, we aim to build models describing how variation in genotype influences variation in phenotype. However, the choice of phenotypes quantitative geneticists choose to analyze is inherently subjective: we typically focus on phenotypes that are practical to measure and/or that are in some sense “important” (e.g. because they are plausibly related to key functions or diseases). These phenotypes are often correlated, presumably because multiple complex traits are often influenced by the same set of core cellular processes. For example, cellular growth rates across a range of different stressful conditions may be determined by a common set of processes such as metabolism, cell wall biosynthesis, DNA repair, and heat or osmotic stress response. This leads to apparent widespread pleiotropy, where individual genetic loci influence many observed phenotypes, presumably because these loci influence one or more core processes that are broadly important across multiple phenotypes.

This perspective suggests that the structure of the correlations between the subjective phenotypes that we choose to measure should contain signatures of the underlying biologically relevant core processes. That is, if we could measure a large and diverse enough set of phenotypes across a sufficiently diverse range of genotypes, the observed phenotypic variation should have a lower-dimensional latent structure that reflects the space of actual core processes. Inferring this lower-dimensional latent structure thus offers the promise of explaining the biological basis of pleiotropy, by identifying the core biological processes and inferring how individual loci influence these core processes to generate the observed phenotypic variation.

Of course, we can only hope to identify core processes which generate variation across the phenotypes we choose to measure, so the core processes we infer will always be limited by this choice. For example, imagine that we measure a set of phenotypes that correspond to the growth rates of yeast cells across a temperature gradient. We might expect that these phenotypes exhibit a correlation structure that reflects three core processes: heat shock response, cold tolerance, and all other temperature-independent factors relevant to the common growth medium. We could then hope to infer the extent to which each genetic locus influences each of the core processes, as well as the mapping between these three core processes and the observed phenotypes. However, if we were to measure additional phenotypes corresponding to growth rates across (for example) different nutrient concentrations, we might find that this splits the temperature-independent core process into additional processes that explain the variation in the new phenotypes.

In this manuscript, we describe how to infer this lower-dimensional latent structure of phenotype space using a penalized matrix decomposition framework [7]. We assume that we have data that describes the map between genotype and some set of measured phenotypes. In general, this genotype-phenotype map can involve nonlinear effects such as interactions between multiple genetic loci (epistasis). However, we focus here on analyzing a standard linear approximation of this map, in which each locus is assumed to have an additive effect on each of the phenotypes, and the observed phenotype is simply a sum of the additive effects of all the relevant loci. This linear map can be represented as an *E* × *L* matrix, **F**, which has columns corresponding to each of the *L* loci and rows corresponding to the effect of these loci on the *E* measured phenotypes. We note that inferring **F** from data on genotypes and corresponding phenotypes can be a complex problem, which we address for one example data set below, but the core of our analysis in this paper assumes that **F** is given and focuses on analyzing the latent structure in this matrix.

In this framework, our problem reduces to inferring lower-dimensional structure in the matrix **F**. While in principle this structure could be nonlinear, we restrict ourselves to inferring a lower-dimensional subspace that can be expressed as a matrix decomposition of **F**. Specifically, we wish to approximate **F** as the product of two matrices, **F** ≈ **WM** + **b**, where **M** is a *K* × *L* matrix that describes the additive effect of each genetic locus on each of *K* putative core processes, and **W** is an *E* × *K* matrix that describes how each core process affects each measured phenotype. In addition, we include a term **b** which represents locus-specific effects on all other processes that contribute equally to the phenotypes measured (and hence cannot be disentangled). For *K < E, L*, this represents an approximation to **F** in terms of a lower-dimensional subspace of *K* core processes. This structure is illustrated in Figure 1a. We emphasize that this decomposition assumes that the map between loci and core processes and the map between core processes and measured phenotypes are both linear, which may not be true in general. We return to this caveat in the Discussion.

**FIG. 1:**
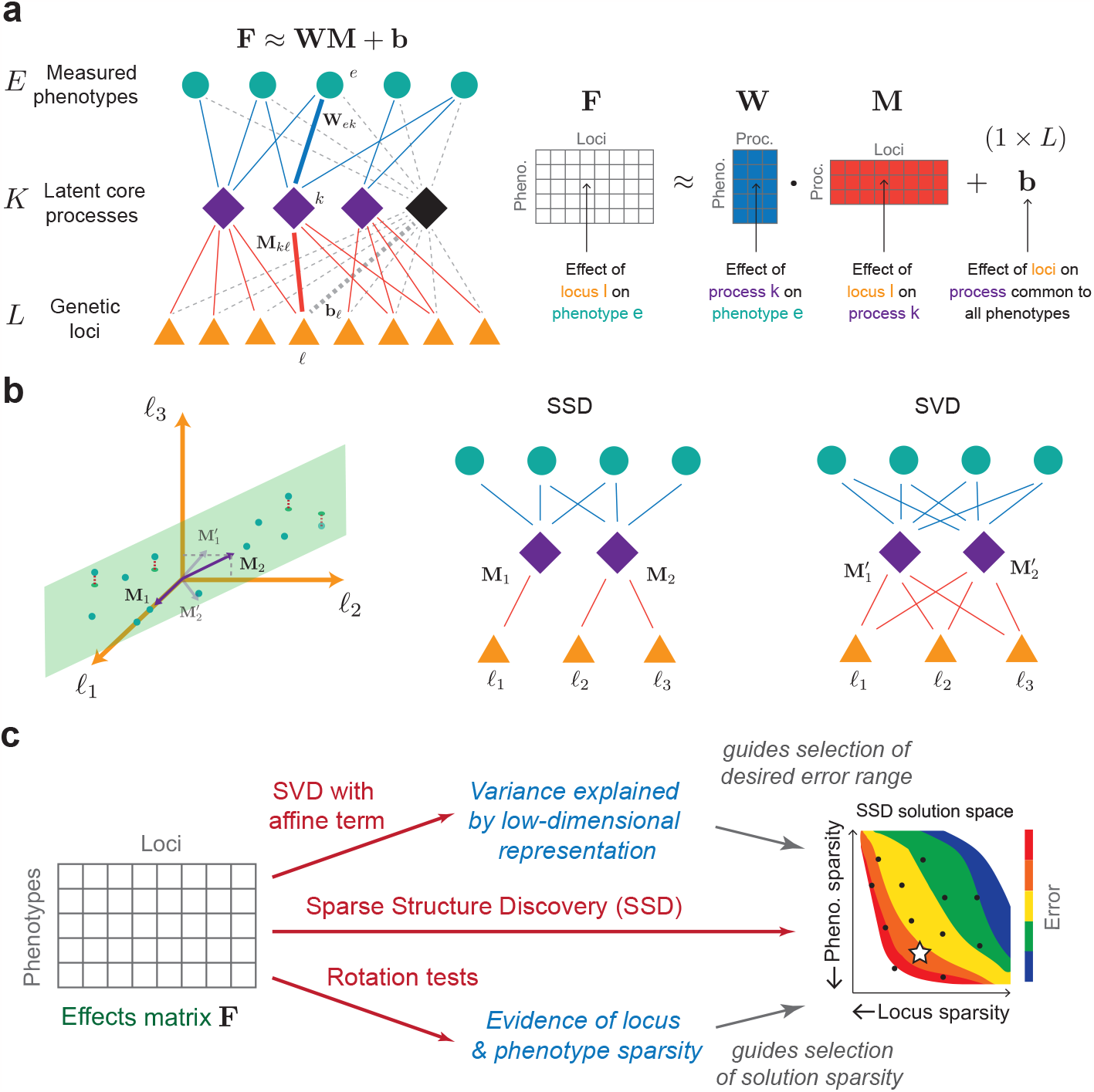
Overview and geometric interpretation of SSD. (a) Sparse structure discovery (SSD) finds a sparse, low-rank approximation for the effects matrix **F** containing the phenotypic effects of *L* loci on *E* phenotypes. (b) Each phenotype (row of **F**) can be viewed as a point in locus-space. The core processes (rows of **M**) can be viewed as vectors that span a lower dimensional subspace, illustrated by the plane. The distances between each phenotype point and the subspace determine the reconstruction error (illustrated by dotted red lines). Since the error is a function of the subspace and there are many matrices **M** which generate the same subspace, many decompositions yield the same error. SSD applied to these phenotypes would favor a sparse decomposition, for example, the core processes **M**_1_, **M**_2_ which here are sparse combinations of (*ℓ*_1_), (*ℓ*_2_, *ℓ*_3_) respectively. SVD applied to the same phenotypes would yield a decomposition with core processes 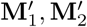 that incur the least error but which are unlikely to be sparse. (c) In our analysis pipeline, we first apply SSD to find a range of decompositions **F** ≈ **WM** + **b** with varying errors and sparsities. The reconstruction error of the SVD solution is used to determine a tolerable error range for SSD solutions. The rotation tests are used to guide the selection of an SSD solution with appropriate levels of sparsity in phenotypes (each phenotype is described by few core processes) and in the loci (each locus is part of few core processes).

Unfortunately, this matrix decomposition problem is underdetermined in general, meaning that for any choice of *K* there are many different pairs of matrices **W** and **M** that approximate **F** equally well. Thus, the fact that a given decomposition gives a good approximation for **F** does not necessarily imply that there is any biological meaning to the core processes inferred. This problem is widely recognized in a variety of fields where matrix decomposition is used to infer lower-dimensional structure in high-dimensional data. To make lower-dimensional structure interpretable, domain-specific knowledge must therefore be used to guide the choice of additional constraints, and optimization algorithms must be developed to efficiently find decompositions that obey the constraints. For example, earlier work has used sparsity [8, 9], non-negativity [10–12] and non-Gaussianity assumptions [13–15] to construct powerful methods for identifying meaningful latent structure in specific contexts where those constraints are appropriate. The success of these approaches motivates our attempt here to find appropriate constraints that enable the efficient and interpretable reconstruction of a lower-dimensional set of core processes from empirical genotype-phenotype maps. Such constraints can be thought of as incorporating a biological “prior” on the features we expect the data to exhibit.

Recently, Kinsler et. al. [16] identified lower-dimensional structure in a dataset describing the effects of a set of yeast mutations on fitness in different environments. Their approach used Singular Value Decomposition (SVD) [17] to find a decomposition with *K < E, L* that approximates the **F** well. However, while SVD finds the *K*-dimensional subspace that explains the most variation for a given *K*, the specific **W** and **M** are selected subject to the constraints that the core processes must be orthogonal and that the first *j* core processes describe the *j*-dimensional subspace that best approximates **F**. It is not clear that these constraints lead to putative core processes with biological meaning. More recently, Pan et. al. [18] introduced an alternative matrix decomposition method, Webster, which is based on regularized dictionary learning [19], and apply it to a dataset describing the fitness of cells exhibiting gene-knockouts in the presence of various genotoxins [20]. This method enforces a hard constraint that each genetic locus affects at most two core processes, which limits the possibility that different loci exhibit different degrees of pleiotropy.

Here, we present a matrix decomposition approach based on the biologically motivated intuition that the lower-dimensional structure of the genotype-phenotype map may be sparse. Specifically, our *Sparse Structure Discovery* (SSD) method finds decompositions where each genetic locus affects a small subset of the core processes (locus-sparsity) and/or each observed phenotype is influenced by a small subset of core processes (phenotype-sparsity). SSD is an example of penalized matrix decomposition, a broad class of matrix decomposition methods that encourage the matrix factors to exhibit particular properties (e.g. sparsity) through hard constraints or regularization [7]. In our case, SSD has separate tunable regularization parameters to control the extent to which locus-sparsity and phenotype-sparsity are encouraged.

The sparsity assumption is consistent with various notions of modularity which have been proposed to explain the evolvability of complex traits [1, 21–25], and with large-scale studies of pairwise gene deletions in yeast, which find that genes cluster together based on their interaction profiles, suggesting their involvement in a small set of common core processes [26]. However, given a matrix **F** representing a genotype-phenotype map, it is not a priori clear the extent to which the data exhibits locus-sparsity and/or phenotype sparsity, making it difficult to justify a specific choice of regularization parameters. To address this, we run our SSD method across a grid of regularization values to obtain an array of decompositions with varying errors, locus-sparsities, and phenotype-sparsities and allow the user to manually select a decomposition of interest for further investigation. Moreover, we have developed two empirical tests to independently validate the extent to which the lower-dimensional structure in an effects matrix **F** exhibits locus-sparsity or phenotype-sparsity, the results of which can then be used to guide the selection of SSD decompositions (Figure 1c). Using these tests, we find evidence of locus-sparsity and phenotype-sparsity across three datasets, motivating the use of these sparsity-enforcing penalties in our SSD method. Further, we show that SSD accurately recovers synthetically-generated maps if at least one of the true **W** or **M** is sparse.

The structure of the paper is as follows. In Section II, we describe the SSD method, explain our empirical tests for sparsity, and demonstrate that SSD accurately recovers core processes in synthetic data. In Section III, we apply our method to three datasets that measure cellular fitness across environments as a function of three different forms of genetic variability. First, we apply SSD to the Kinsler et. al. dataset [16] describing fitness effects of adaptive mutations identified during a laboratory yeast evolution experiment and compare SSD to the SVD-based analysis presented in [16]. Second, we apply SSD to data describing how single gene knockouts in human cell lines affect fitness in the presence of genotoxic agents [20]. We find that, compared to the Webster analysis of the same dataset [18], SSD solutions exhibit lower error with comparable average sparsity, a more interpretable process-phenotypes map, and a broad range of pleiotropy across loci. Third, we analyze a large-scale quantitative trait locus (QTL) mapping experiment [27], which measured 18 growth rate phenotypes in about 100,000 F1 offspring of a cross between two related budding yeast strains. For this data, we first develop a joint mapping approach to arrive at an additive effects matrix **F**, which we do using a pipeline based on *ℓ*_2,1_-penalized regression (see SI).

## II. SPARSE STRUCTURE DISCOVERY

As described above, our method assumes we begin with an empirical linear genotype-phenotype map, represented as an *E* × *L* matrix **F** which describes the additive effect of each of the *L* genetic loci on each of the *E* measured phenotypes. Our goal is to find latent structure in this genotype-phenotype map of the form **F** ≈ **WM** + **b**. Note that since we will generally assume that *K < E, L*, the matrices **W** and **M** contain fewer total parameters than **F** (i.e. this is a simpler description of the data). Thus, this factorization will in general only be an approximation, both because there is presumably error in the estimation of **F** and because the division into *K* core processes is a simplifying assumption that will inevitably neglect some aspects of the full complexity underlying each measured phenotype.

Given that the factorization of **F** is approximate, a natural goal would be to find matrices **W** and **M** that minimize the error in this approximation. This is the motivation underlying singular value decomposition (SVD), which finds a factorization of **F** that minimizes the squared Frobenius reconstruction error (i.e. lowest squared error 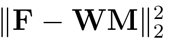). However, this error minimization alone is not sufficient to uniquely determine the factorization. Instead, any factorization that describes the same lower-dimensional subspace will perform equally well, as illustrated in Figure 1b. This is a general problem: for any set of core processes, represented by the rows of **M**, that achieve a given reconstruction error, there are infinitely many sets of other processes that achieve the same error (obtained by changing the basis of the subspace, e.g. by rotating the rows of **M** in the subspace they generate). SVD chooses a particular unique solution to resolve this degeneracy by defining the first core process to be the one-dimensional subspace that minimizes the error for *K* = 1, the second core process to be orthogonal to the first and minimize the error for *K* = 2, the third to be orthogonal to the first two and minimize the error for *K* = 3, and so on. While this is a reasonable and well-defined procedure, there is no reason to believe that the core processes defined in this way will be biologically meaningful.

Here we define an alternative method for matrix decomposition. Like SVD, our approach attempts to minimize the Frobenius reconstruction error. However, we add two additional constraints based on *sparsity*. Specifically, we aim to find a locus to core process map **M** in which each locus participates in only a few processes (i.e. most entries in this matrix are 0). We refer to this as locus-sparsity. Analogously, we aim to find a core process to phenotype map **W** in which each phenotype is affected by only a few core processes (i.e. most entries in this matrix are also 0). We refer to this as phenotype-sparsity.

We do not necessarily assume that both types of sparsity exist in a given dataset. Instead, our framework allows us to impose constraints on either or both types with a tunable stringency (and below we describe how the choice of this stringency can be guided by empirical validation tests). To be precise, our *Sparse Structure Discovery* (SSD) method aims to find the matrix decomposition **F** ≈ **WM** + **b** that minimizes

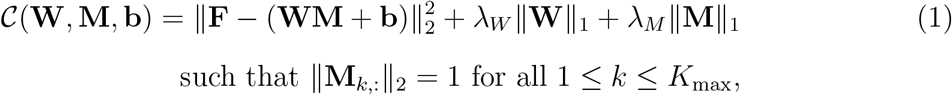

where 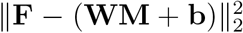 is the squared Frobenius error, ∥**W**∥_1_ is an *ℓ*_1_-norm measure of the phenotype-sparsity, and ∥**M**∥_1_ is an *ℓ*_1_-norm measure of the locus-sparsity. Equation 2 is a variant of the penalized matrix decomposition formulations studied in [7]. The parameters *λ*_*W*_ and *λ*_*M*_ determine the relative weighting of the accuracy, phenotype-sparsity, and locus-sparsity objectives (higher *λ*_*W*_ will yield solutions that are more phenotype-sparse, and higher *λ*_*M*_ will yield solutions that are more locus-sparse). We note that when these regularization parameters *λ*_*W*_ and *λ*_*M*_ are sufficiently large, the method will assign no loci to some of the core processes, thereby automatically picking a number of core processes *K* smaller than the input upper bound *K*_max_. We include the constraint requiring that the Euclidean norm of each core process is one to ensure that the core processes are all of the same scale; the magnitude of the vectors of **W** may vary, reflecting that some phenotypes are more sensitive to the core processes than others.

For fixed values *λ*_*W*_, *λ*_*M*_, and *K*_max_, SSD will yield a unique set of **W, M** and **b**. However, a key challenge is to choose values of these parameters to determine an appropriate weighting of the accuracy, phenotype-sparsity, and locus-sparsity objectives that will produce a decomposition with plausible biological meaning. To do so, we first apply our method for a range of values *λ*_*W*_ and *λ*_*M*_ to produce a variety of decompositions that vary in reconstruction error, number of core processes, locus-sparsity, and phenotype-sparsity. In every case, the SSD decomposition will have higher reconstruction error than the SVD decomposition with the same number of processes because of the additional constraints. We therefore use the SVD error as a guide to select a desired reconstruction error range, and select sparse decompositions of interest that fall within this range. The choice between these can then be guided by the empirical test described below, which we developed to determine the extent to which an input matrix **F** exhibits a low-dimensional structure with locus-sparsity or phenotype-sparsity. Figure 1c illustrates the pipeline.

### A. Empirical validation of sparsity constraints using rotation tests

To validate our choice of sparsity assumptions, we designed heuristic tests to determine whether a given dataset **F** exhibits signatures of locus-sparsity or phenotype-sparsity. We do not assume that the linear term **b**, which describes the effects of loci on processes that do not vary across the phenotypes, is necessarily sparse. For the purposes of this test, we therefore first subtract the mean effect across phenotypes for each locus from **F**, as an approximation of **b**. To test for locus-sparsity, we then apply a random orthogonal matrix **O** to the empirical genotype-phenotype map **F** to produce a matrix **F**^′^ = **FO**. This rotation conserves low-dimensional structure in **F** and leads to the same SVD error but disrupts any potential locus-sparsity. We then apply our SSD method with a range of weights on the locus-sparsity objective to obtain a range of decompositions for **F** and **F**^′^ that exhibit varying locus-sparsities and reconstruction errors. If the input matrix **F** truly has locus-sparsity, our method will consistently find sparser solutions for **F** than for **F**^′^ across a range of reconstruction errors. If so, we consider this to be evidence of locus-sparsity in **F**.

To gain intuition for this test, consider an example with five loci and two core processes, with loci *ℓ*_1_ and *ℓ*_2_ both affecting core process 1 (with equal weight) and loci *ℓ*_3_, *ℓ*_4_ and *ℓ*_5_ all affecting core process 2 (also with equal weight). The rows of the matrix **F** will each have the form (*α, α, β, β, β*), where *α* and *β* describe the effect of the first and second processes on the phenotype corresponding to that row, respectively. In other words, the phenotype values lie on a 2D plane in 5D space. This plane contains the sparse vectors (1, 1, 0, 0, 0) and (0, 0, 1, 1, 1), which describe the two core processes, and every point on the plane can be written as a weighted sum of these vectors. Now, imagine that we randomly rotate **F**, producing a matrix **F**^′^ which has rows that lie on a rotation of the 2D plane containing the rows of **F** and columns that correspond to random linear combinations of the actual genetic loci. Since the rotation was random, the 2D plane containing the rows of **F**^′^ is a random 2D plane in 5D. Most 2D planes in 5D are not spanned by two sparse basis vectors. Therefore, while it is still possible to find two vectors such that each row of **F**^′^ can be written as the weighted sum of these vectors (the low-dimensional structure is preserved), the two vectors almost certainly will not be sparse.

To test for phenotype-sparsity, we use an analogous method, except that we rotate the columns of **F** to obtain **F**^′^ = **OF** and vary the phenotype-sparsity objective in SSD to test whether SSD consistently finds sparser solutions for **F** than **F**^′^ across a range of reconstruction errors.

### B. Sparse structure recovery on synthetic data

To validate our method, we constructed synthetic genotype-phenotype maps with lower-dimensional latent structure of varying sparsity. That is, for a given *E, L*, and *K*, we construct simulated data matrices **F** = **WM** + *η* by randomly choosing **M** and **W** as described below. The noise *η* in each element is drawn independently with scale 0.3 times the standard deviation of the entries in **WM**. We construct simulated **F** matrices across a range of sparsities in **M** and **W**. Specifically, for **M**-sparsity *p*, entries are non-zero with probability *p*, and if non-zero, the entry is drawn from a standard normal. We then normalize **M** so that each row is a unit vector. We generate **W** analogously with **W**-sparsity *q*, but without normalization. By not normalizing **W**, we allow for the possibility that some phenotypes are influenced more strongly by the core processes than others.

We begin by constructing four sets of simulated data: one with both locus-sparsity and phenotype-sparsity (*p* = 0.2, *q* = 0.2), one each with only one type of sparsity (*p* = 0.2, *q* = 1 and *p* = 1, *q* = 0.2), and one with neither (*p* = 1, *q* = 1). For each set, we first applied the locus and phenotype rotation tests. The results are presented in the left column of Figure 2. Note that the presence of the gap between the error curves for **F** and rotated **F**^′^ in the locus (phenotype) rotation test depends on whether **M** (**W**) is sparse. Repeating this test across a range of locus and phenotype sparsities, we show that the size of the gap grows continuously with sparsity (Figure S1).

**FIG. 2:**
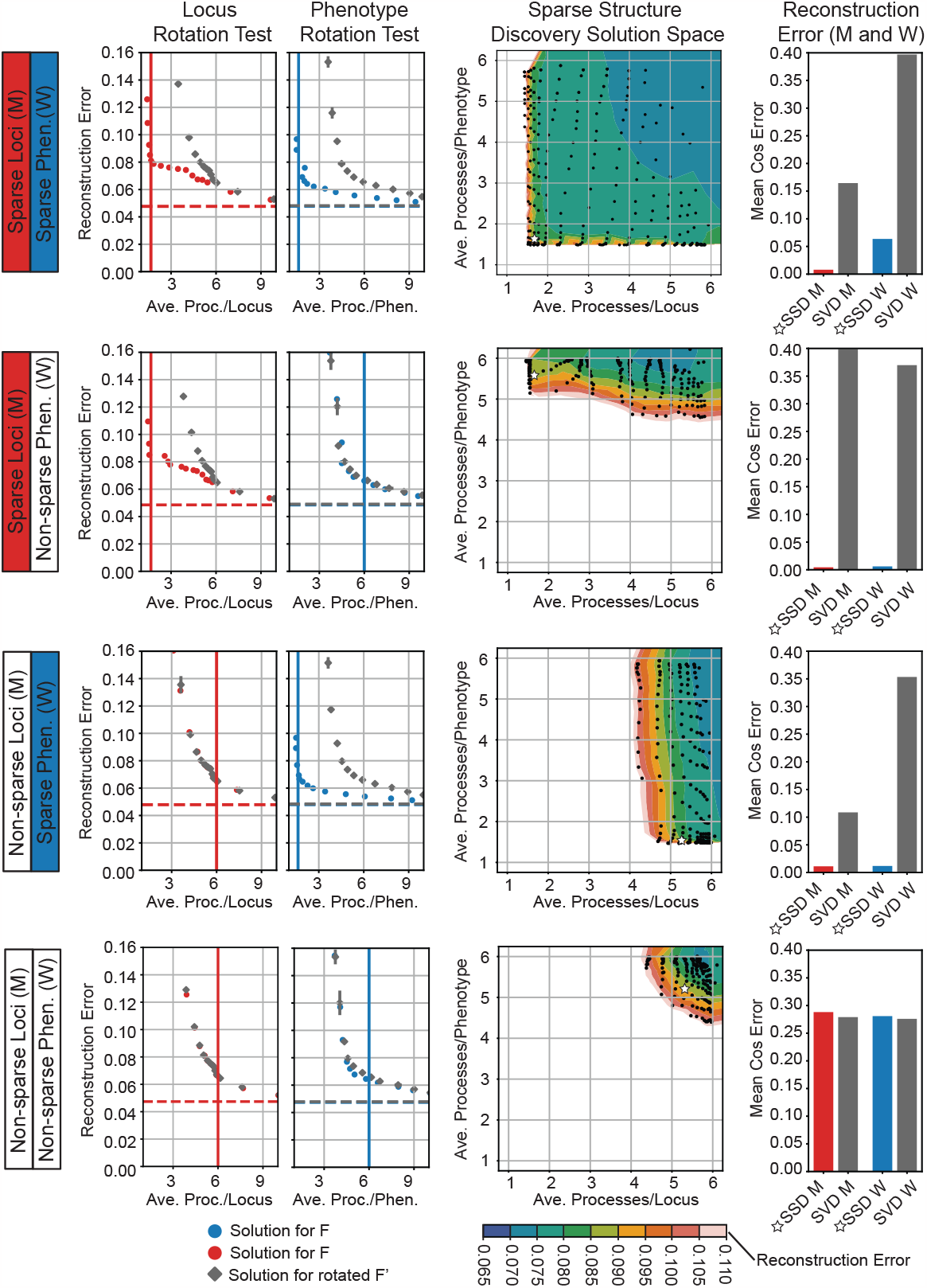
SSD on synthetic data. Each row corresponds to a synthetic additive effects matrix *F* = **WM** + *η* generated with different sparsities in **M** and **W**. All examples have *E* = 96 phenotypes, *L* = 200 loci and *K* = 6 true core processes. First column: Locus (phenotype) rotation test illustrates that when **F** is generated with sparsity in **M** (**W**), there is a gap between the sparsity of the SSD solutions for **F** in blue (red) and rotated **F**^′^ = **FO** (**F**^′^ = **OF**) in grey. Each scatter point corresponds to a solution for a particular value of the regularization parameter *λ*_*M*_ (*λ*_*W*_). Error bars are over three random rotations; error bars were often so small that they were not visible over the scatter point. The red and blue horizontal dashed lines indicate the 6-component SVD reconstruction error. The grey dashed line indicates the average 6-component SVD reconstruction error of **OF**. Note that the SVD error for **FO** is equal to that for **F**. The vertical red (blue) line indicates the average processes per locus (phenotype) for the true **M** (**W**). Second column: Each scatter point depicts the average processes per locus/phenotype of an SSD solution. The colored background illustrates the interpolated reconstruction errors of the solutions. The solutions selected for further investigation are marked by a star. Third column: The mean cosine error between each row of the inferred **M** (column of inferred **W**) for the selected SSD solution and for the 6-component SVD solution and the true **M** (**W**). The error in **W** in the first row is almost exclusively due to 4 phenotypes that use no processes, but are assigned very small weights in some processes by SSD.

Next, we evaluated whether SSD can accurately reconstruct the true **M** and **W** matrices. We applied our SSD method to each dataset across a range of locus-sparsity (*λ*_*M*_) and phenotype-sparsity (*λ*_*W*_) constraints and selected one decomposition using the SVD error and rotation tests as guides. The reconstruction error of the SVD decomposition on each dataset is in the range 0.047 −0.049. Keeping in mind that any SSD solution will necessarily have higher error, we focus on “low-error” decompositions with error up to 0.85, illustrated by dark green, teal, and blue in the space of SSD decompositions (Figure 2, center column). When either **M** or **W** is sparse, the space of solutions below a certain error is rectangular (Figure 2). In this case, we select a decomposition that exhibits the most sparsity for the chosen error criterion, that is, the bottom left vertex of the rectangular contour (indicated by a white star in Figure 2). When both **M** and **W** are not sparse, we pick a solution with similar degrees of locus and phenotype sparsity that satisfies our error criterion.

Finally, we compared the **M** and **W** of the selected SSD solutions to the true **M** and **W** matrices using a cosine error metric described in the Methods (third column of Figure 2). We find that exhibiting sparsity in either **W** or **M** (first three rows) suffices for SSD to accurately reconstruct both **W** and **M**. Given a non-redundant set of core processes **M**, there is a unique set of phenotype weights **W** that best reconstruct **F** (and vice-versa for **W**). In contrast to SSD, the SVD decompositions are unable to accurately reconstruct **M** and **W**, despite lower reconstruction errors when reconstructing **F**.

The phenotypes constructed as described in this section are correlated in so far as each is a random linear combination of a common set of core processes. However, empirical studies may measure phenotypes with non-trivial structure, e.g. fitness measurements where the same environmental perturbations are added to various growth mediums. To validate the rotation tests and SSD in such a setting, we generated synthetic data with a hub-and-spoke structure. Specifically, we introduce “hub” phenotypes (representing the growth mediums) whose effects are a random linear combination of a common set of core processes and “spoke” phenotypes (each representing a growth medium with a perturbation) whose effects are a linear combination of the corresponding hub phenotype and one core process representing the perturbation (Figure S2a). See SI for further details.

Next, we apply the rotation tests to the hub-and-spoke synthetic data and find evidence of both locus-sparsity and phenotype-sparsity (Figure S3). We find that a selected SSD solution exhibiting both types of sparsity accurately recovers the initially described generative structure. If we instead ignore evidence of locus-sparsity and select an SSD solution that exhibits a greater degree of phenotype-sparsity and little locus-sparsity, the decomposition resembles an alternate generative structure where each hub phenotype is instead described by a single core process (Figure S2b). In contrast, SVD finds a solution with lower reconstruction error but with matrices **M** and **W** that lack any clear relationship to the core processes that generated the synthetic data.

## III. APPLICATIONS TO EMPIRICAL DATA

### A. Fitness effects of adaptive mutations in yeast

To illustrate the applicability of our framework, we first analyze data from a recent study by Kinsler et. al. [16]. This study attempted to infer a lower-dimensional latent structure of phenotype space by measuring the fitness effects of a set of specific yeast mutations across a range of environmental perturbations. Specifically, they isolated 292 yeast strains from an earlier laboratory evolution experiment, each of which contains one or a few putatively adaptive mutations. They measured the fitness of each of these strains across a set of 45 environments (Figure 3a). Based on these measurements, they divided the 45 environments into 25 “subtle” perturbations (in which fitness effects of mutations vary only slightly) and 20 “strong” perturbations. Applying SVD on the data from the subtle perturbations, they identified an eight-dimensional subspace that explains most of the variation in the data across these perturbations. They then showed that this latent structure can also predict the fitness effects of the mutations across the 20 “strong” perturbations, which they interpret as evidence that the subtle perturbations reveal a “local” modularity that is able to predict the global pleiotropic effects of adaptation in this system.

**FIG. 3:**
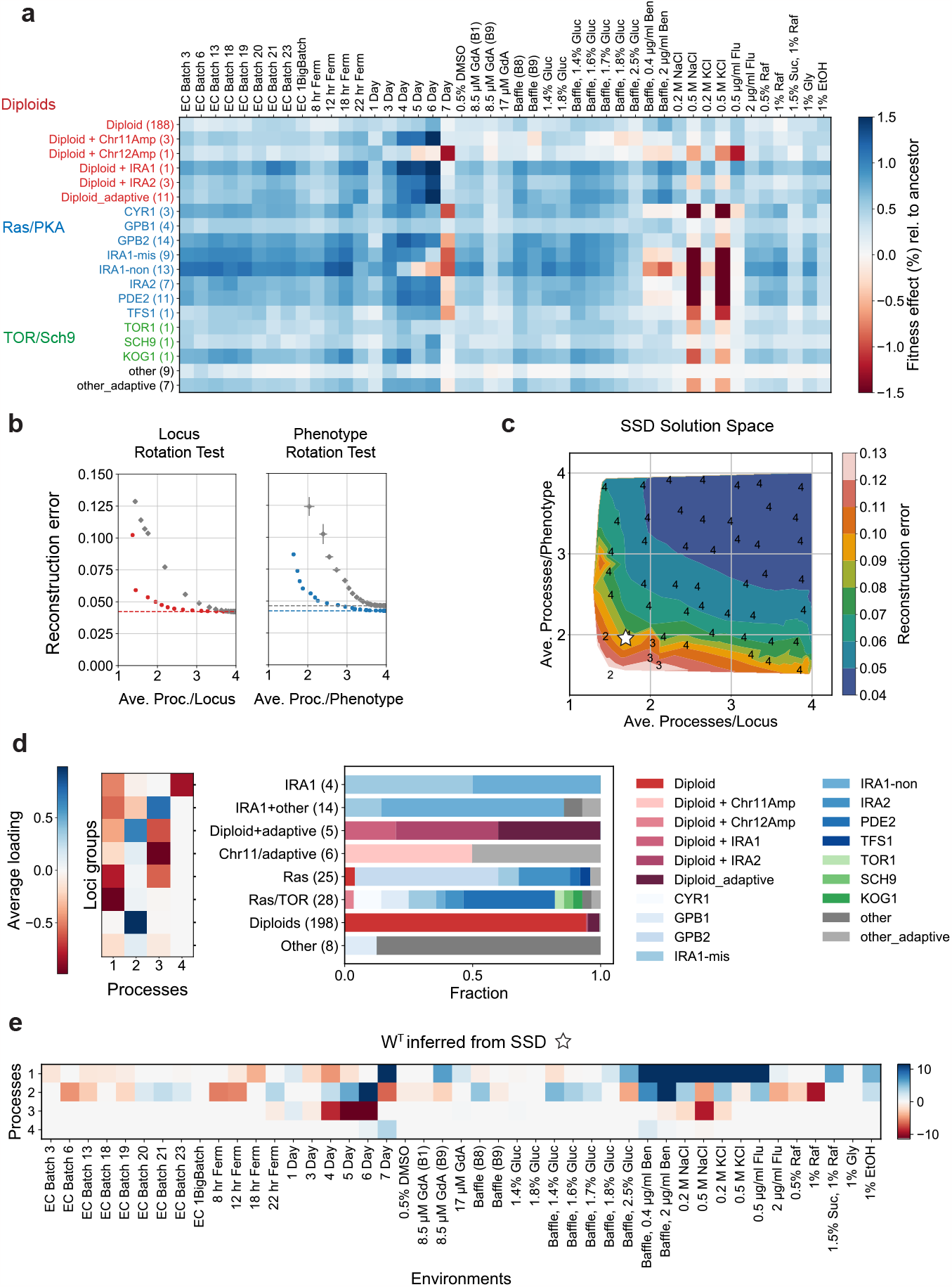
SSD applied to pleiotropic fitness effects of adaptive mutations in yeast. (a) A reduced representation of the effects matrix **F** (45(*E*) × 288(*L*)) where the effects of mutations with common annotations are grouped together. The number of mutations with each annotation is shown in parenthesis. (b) The locus and phenotype rotation tests show extensive sparsity in both the process-phenotype and locus-process maps. (c) The solution space illustrating highly sparse solutions with low reconstruction error. The integers indicate the number of processes in the solution. The chosen solution with 8.5% error is marked with a white star. (d) The **M** matrix with loci clustered into 8 groups based on linkage clustering of loci with a modified cosine similarity metric (see Methods). On the right, the fraction of loci types in each of the 8 groups is shown. The number of loci in each group is shown in parenthesis next to its label. (e) The process-phenotype map **W**.

We sought to investigate whether our SSD method can recover an alternative sparse lower-dimensional structure in the Kinsler *et. al*. data. Rather than divide environments into “subtle” and “strong” perturbations, we took the entire mutational effects matrix representing 288 strains across 45 environments as our input **F** (we use 288 instead of the original 292 due to a minor difference in a pre-processing step, see SI). We then applied our locus and phenotype rotation tests (Figure 3b), which confirm that there is strong evidence for sparsity in both the process-phenotype map (**W**) and the locus-process map (**M**). Note however that removing most diploids from this data (one key type of mutation that represents 188 of the 288 mutations studied) eliminates sparsity in **M** but not in **W** (Figure S4). Further analysis (discussed below) finds that the diploids predominantly affect one core process and thus the locus-sparsity indicated by the rotation test can be explained by the large number of diploids in the data. This is not an issue for applying SSD, as SSD requires sparsity in only one of **W** and **M**.

We find that SSD can identify a sparse, 4-dimensional approximation of **F** that incurs less than 8% error in reconstructing the original **F** (Figure 3c). For concreteness, we focus here on the sparse solution indicated by the white star in Figure 3c, which has four core processes and an average sparsity of about 1.5 processes per locus and 2 processes per environment. In Figure S5, we highlight the differences between the SVD and SSD solutions. By construction, the SSD solution has a higher reconstruction error than the corresponding SVD solution (7.5% error for the sparse SSD solution, compared to 4% error for the 4-dimensional SVD solution). We find that the SVD solution on a training set also shows lower error in predicting the fitness effects in held-out environments (the 20 strong perturbations or a random subset of 9 environments) compared to the SSD solutions of equal rank (Figure S5). This suggests that SVD tends to find a better low-rank approximation, even when it fails to find meaningful (and potentially sparse) basis vectors (see Discussion). To highlight this point, if SVD finds the locus-process and process-phenotype maps **M**_SVD_, **W**_SVD_ on the training set, it can be mathematically shown that the maps **M**^′^ = **OM**_SVD_, **W**^′^ = **W**_SVD_**O**^*T*^ for any arbitrary orthogonal matrix **O** will match SVD’s generalization error. In contrast, the SSD solution is significantly sparser than the SVD solution (Figure S5) at the expense of a larger generalization error. Thus, while SVD by construction finds the subspace with the lowest reconstruction error, the SSD approach is able to identify sparse basis vectors, capturing the sparsity in the genotype-phenotype map suggested by the rotation tests.

To examine if loci with similar effects on core processes identified by SSD align with existing annotations, we further clustered loci into eight groups by comparing the columns of the **M** matrix with a modified cosine metric (Methods). We observe that core process 1 is enriched for mutations in genes involved in the Ras and TOR pathways (Figure 3d). Missense and nonsense mutations in IRA1 (also involved in the Ras pathway) clustered in the “IRA1+other” group have additional pleiotropic effects on core process 3, which has a large influence on fitness in environments with an extended stationary phase (4,5 and 6 Day environments in Figure 3e). Diploids are primarily enriched in core process 2, which has broad pleiotropic effects across environments. Diploids with additional mutations in IRA1/2 (clustered in the “Diploid + adaptive” group) exhibit effects that combine the effects shown independently by IRA1/2 in the Ras cluster and the Diploids cluster. Thus, the core processes identified by SSD do appear to have some correspondence with our prior expectations. To ensure that the many diploids do not significantly bias our results, we repeated this analysis on a reduced dataset which excludes a random subset of 168 of the 188 diploids, finding similar features in the **W** and **M** maps despite lower average sparsity in **M** (Figure S4).

Finally, it is easier to read off hypotheses from a sparse SSD decomposition than from a dense SVD decomposition (Figure S5b). For example, since SSD core process 3 almost exclusively impacts environments with an extended stationary phase (4,5 and 6 Day), it is reasonable to hypothesize that loci involved in this core process influence a pathway relevant in stationary phase. In contrast, each SVD core process affects most environments (Figure S5c), thereby confounding an analogous interpretation. The SSD solution further suggests that diploidy primarily contributes to core process 2, and the contribution of this process across environments is a succinct summary of its effect. For the SVD solution, the diploids do not form a single cluster (Figure S5c,d), and no such summary is apparent.

### B. Robustness of gene knockouts to genotoxins in human cell lines

Next, we apply our SSD method to the genotoxic fitness screen collected in [20] and curated in [18] (Figure 4a). This dataset was constructed by performing CRISPR-Cas9 knockouts on an immortalized human cell line (RPE1-hTERT) and subjecting each knockout variant to 31 genotoxic stressors. We show that the core processes described by our SSD decomposition are enriched for particular gene annotations and compare our decomposition to one identified by Webster [18].

**FIG. 4:**
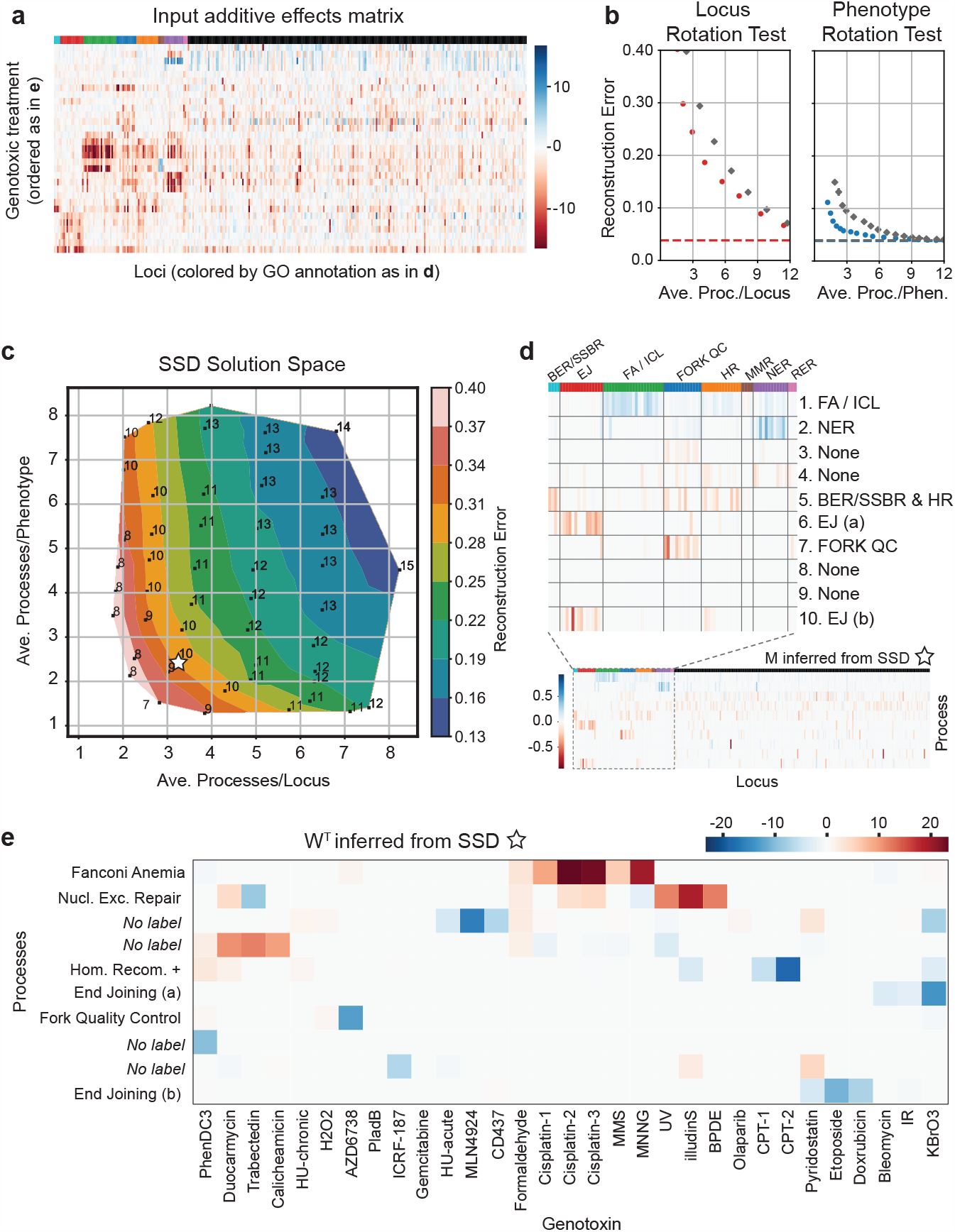
SSD applied to dataset of human cell responses to gene knockouts under genotoxic stressors. (a) The input additive effects matrix **F** generated by [20] and curated by [18]. (b) The locus and phenotype rotation test indicate there is both locus and phenotype sparsity. (c) The space of solutions found by SSD. The integers indicate the number of processes in the solution. The white star indicates the solution that we illustrate in (d) and (e). (d) Sorting the loci by GO annotation in the locus-process map **M** reveals that certain processes are enriched for particular annotated functions. (e) The process-phenotype map **W** demonstrates that the response to each genotoxin can be explained by a small number of core processes.

Our rotation tests find evidence of both locus and phenotype sparsity in this genotoxin data (Figure 4b). Phenotype-sparsity is not assumed by Webster [18], suggesting that SSD may lead to a more interpretable process-phenotype map. In order to compare directly to the Webster decomposition analyzed in [18], we restrict our attention to SSD solutions that have the same number of core processes (*K* = 10). Note that alternate SSD solutions with fewer core processes incur more error. Guided by the results of the rotation tests, we select a solution that is sparse in both loci and phenotypes (3.3 average-processes-per-locus, 2.5 average-processes-per-genotoxin), indicated by the white star in Figure 4c.

First, we evaluate whether the core processes described by our solution are enriched for loci with particular functional effects. We organize the locus-process map **M** by the loci annotations compiled in [20] and observe that core processes 1, 2, and 7 are enriched for loci involved with the the repair of interstrand cross-links (ICLs) by Fanconi Anemia (FA) proteins, nucleotide excision repair (NER), and DNA replication fork quality control (FORK QC) respectively (Fig 4d). Loci involved with end joining are primarily split between core processes 6 and 10. Finally, core process 5 is enriched for loci involved with base excision repair (BER) and single-strand break repair (SSBR) as well as homologous recombination (HR). The functional meaning of the other four processs are not immediately clear from the annotations so we leave them unlabeled; investigating the loci with the strongest effects could elucidate their meaning, as was done by Pan *et. al*. [18]. Figure 4e illustrates the process-genotoxin map **W**; the sparsity indicates that a small number of core processes explain the effect of each genotoxic stressor.

In the SI, we further describe the differences between Webster and SSD and compare the decompositions of this dataset found by each method. Our SSD method more accurately reconstructs the the additive effects matrix while exhibiting more phenotype-sparsity and only slightly less locus-sparsity. Moreover, our SSD decomposition exhibits variation in the degree of pleiotropy across loci, measured by the number of processes each locus participates in (Figure S6).

### C. The genotype-phenotype map of a yeast cross

Next, we analyze data from a recent study [27] analyzing genotypes and phenotypes of *N* ≈ 100, 000 F1 haploid yeast offspring (segregants) of a cross between RM (a European wine strain) and BY (a standard lab strain). These two parental strains differ by *S* ≈ 42, 000 single-nucleotide-polymorphisms (SNPs), leading to a highly diverse set of genotypes in the segregant pool. This earlier work measured the fitness (growth rate relative to the parental BY strain) of each of the segregants in *E* = 18 environments using a bulk barcode-based phenotyping assay.

The base condition for most of these environments is propagation in batch culture with 1:128 dilutions every 24 hours in rich laboratory media (YPD) at optimal temperature (30C). We refer to this as the 30°C environment. Other environments are then constructed by adding stressors to this base condition (e.g. lithium, 4-nitroquinoline oxide, ethanol), by varying the temperature (23°C to 37°C), by using defined media with various carbon sources (glucose, mannose, raffinose) instead of YPD, and by using complex natural media (molasses).

To apply SSD to this data, we must first infer the genotype-phenotype map for each of these 18 environments (i.e. we must infer **F**). This is a complex problem; Ba *et. al*. [27] includes an extensive discussion of the challenges associated with this inference and introduces a modified stepwise forward search procedure for this purpose. A particular difficulty is that this mapping is typically not able to precisely pinpoint specific loci that affect each phenotype. Because our goal is to use the genotype-phenotype map across these different environments to infer lower-dimensional latent structure, we adopt a simpler approach here. Instead of identifying putative causal loci separately for each phenotype, we use a penalized regression approach to jointly identify a sparse set of loci that explain the fitness across environments (see SI). Then, we use a statistical test to establish a confidence interval for the location of each putative causal locus. This procedure identifies 1089 genomic regions containing putative loci and their fitness effects in the 18 environments. We use this 18 × 1089 matrix as the effects matrix **F** for SSD, represented schematically in Figure 5a.

**FIG. 5:**
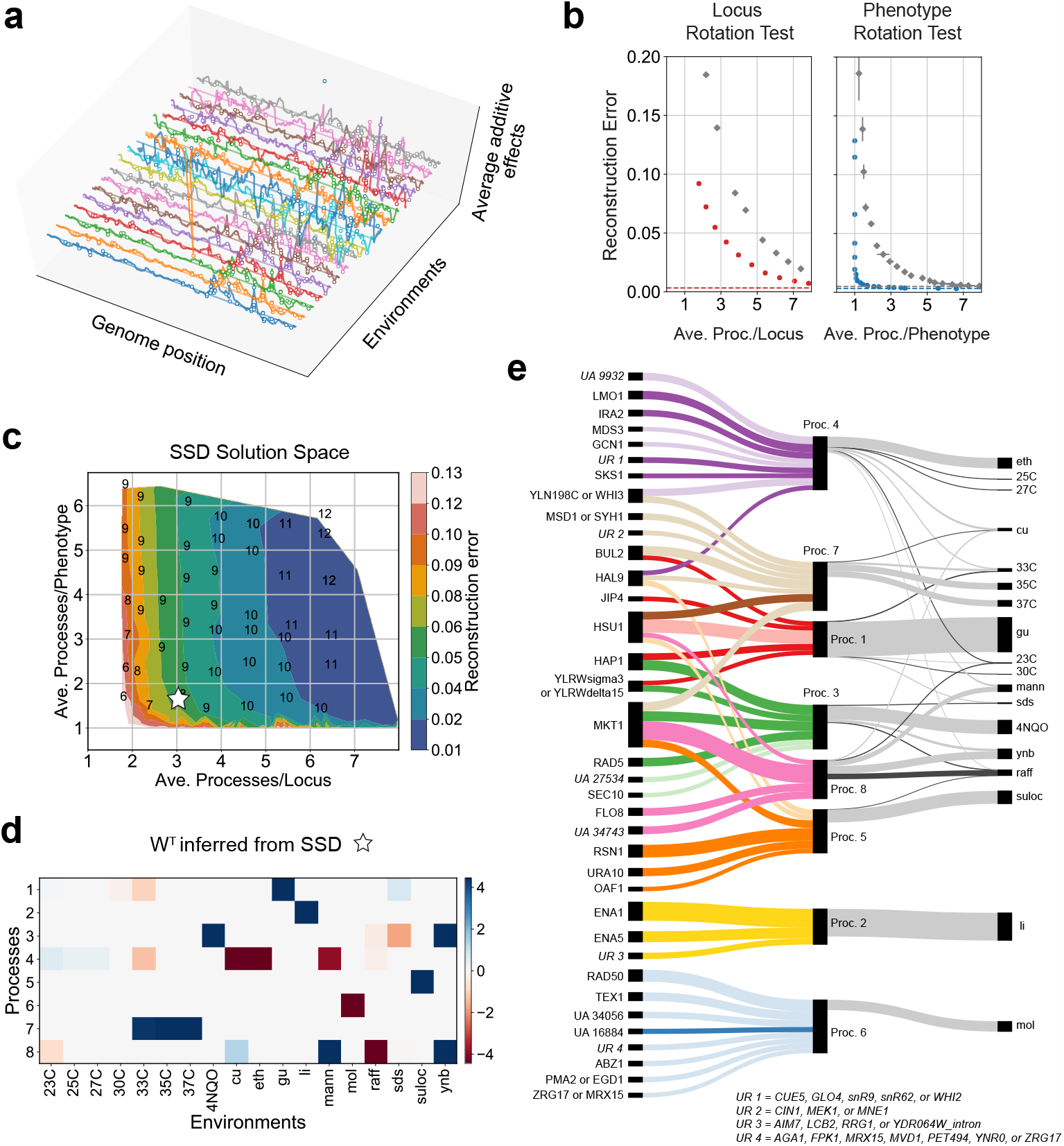
SSD on genotype-phenotype data from a yeast cross. (a) The average additive effects of *S* ≈ 42, 000 genetic loci, estimated using unpenalized linear regression for each of the 18 environments independently. The environments are arranged from bottom to top as arranged in panel d from left to right. Note that the correlations in average additive effects across neighboring loci due to linkage. (b) The loci and phenotype rotation tests, showing extensive sparsity in the process-phenotype map and moderate sparsity in the locus-process map. (c) The solution space has a landscape reflecting the sparsity in the process-phenotype map. The integers indicate the number of processes in the solution. The solution picked for downstream analysis is starred. (d) The process-phenotype map, **W**. (e) A Sankey figure illustrating the locus-process map **M** for the large effect loci in each core process (color) and the process-phenotype map **W** (grey). The width of each line is proportional to the magnitude of the value in **M** or **W**. In **M**, the lighter (darker) shade of each color indicates that the RM (BY) allele contributes positively to the process. In **W**, light and dark grey indicate positive and negative contributions to the phenotypic measurement respectively. The signs of the core processes are adjusted so that they impact most phenotypic values positively.

We next apply the loci and phenotype rotation tests (Figure 5b), finding evidence for extensive sparsity in the process-phenotype map **W** and moderate levels of sparsity in the locus-processes map **M**. The SSD solution space shows an error landscape that favors low-rank (*K* ≈ 6 − 9) approximations to **F** which are sparse in **W** (Figure 5c). We focus here on the *K* = 8 solution indicated by the white star in Figure 5c, which represents a trade-off between achieving high sparsity in **W** and moderate sparsity in **M** while retaining relatively low reconstruction error. We verified that this solution explains a fraction of variance on a test set of genotypes comparable to that explained by the full **F** and the 8-component SVD solution (Figure S7a). Other reasonable choices of solutions lead to qualitatively similar results (Figure S7b).

In Figure 5d we show the resulting inferred **W**. We find that this matrix is sparse and has some intuitive features. First, we note that the term **b** in our SSD decomposition represents a constant effect of each locus on all of the measured phenotypes (i.e. the aspect of the genotype-phenotype map that is constant across all the environments). The inferred **W** then represents how the loci in a given process produce deviations from these constant effects across the different environments. We find that none of the inferred processes have substantial weight in **W** for our 30°C environment, indicating that **b** fully captures the genotype-phenotype map for this environment. This is intuitive, given that this environment is the basis for all other conditions. The environments which represent this same condition at slightly lower temperatures are also largely captured by **b**, though processes 4 and 8 do become slightly more important as we decrease the temperature. As we increase temperature, we find that process 7 becomes important, suggesting that this process is associated with high temperature response. Several processes are specific to given environments (e.g. process 1 primarily affects fitness in guanidinium chloride (gu), process 2 affects fitness in lithium (li), process 5 in suloctidil (suloc), and 6 in molasses (mol)). Some of these processes, such as processes 2 and 6, contain a largely non-overlapping set of loci that affect their respective environments (li and mol) in addition to the constant effects captured by **b**. Finally, processes 3, 4, and 8 reflect processes that influence a few conditions, including some observed trade-offs (e.g. between fitness in raffinose and ynb or mannose).

In SI Table 1, we provide a list of the ORFs localized to each putative causal locus, GO annotations and descriptions from the Saccharomyces Genome Database [28], and their influence on each core process (i.e. value in **M**). In Figure 5e we show a Sankey figure that illustrates **W** and the most prominent features of **M**. This figure shows both how a number of key loci affect each of the processes (i.e. features of **M**), and how these processes in turn affect fitness in each of the environments (i.e. **W**). For example, we see that the genes ENA1 and ENA5 are the primary contributions to process 2, and that this process primarily influences fitness in lithium. This is consistent with prior expectations, as the ENA cluster is involved in salt tolerance and is known to be important for lithium tolerance [29]. Similarly, we see that BUL2, known to affect heat-shock element mediated gene expression (see SI Table 1), is the primary contributor to process 7, which influences fitness in the high temperature environments. In addition, some loci which are known to have large effects on fitness across these conditions (e.g. MKT1, IRA2) are also represented in **M**. There are also many other loci (some of unknown function and other unannotated genes) that play a role, and the rationale for these patterns is unclear. Additional experiments measuring fitness across a larger set of environments may help further disentangle structure in this genotype-phenotype map, and help resolve additional processes.

## IV. DISCUSSION

Extensive work in quantitative genetics has aimed to develop models that explain the relationship between genotype and a variety of different phenotypes. This work often finds widespread pleiotropy, where specific genetic variants affect multiple phenotypes, creating a complex pattern of correlations between phenotypes. Using these patterns to infer a lower-dimensional structure in the map between genotype and multiple phenotypes is an important goal, which offers the promise of identifying a biologically meaningful explanation for observed patterns of pleiotropy.

A central challenge in achieving this goal is that discovering lower-dimensional structure in high-dimensional data is fundamentally underdetermined. Thus, we must always choose some set of objective functions and/or constraints as the basis for any such decomposition. This choice is inherently somewhat arbitrary, and it is not immediately clear how to select objectives and constraints that will lead to solutions that reflect biologically meaningful structure in the data.

In this paper, we address this challenge using a penalized matrix decomposition framework, *Sparse Structure Discovery* (SSD), which allows us to identify a low-dimensional set of “core processes” that concisely explains the observed patterns of pleiotropy in genotype-phenotype data. The method uses sparsity as a key constraint to decompose a model for how genotype influences multiple phenotypes into two linear sparse lower-dimensional maps: a map between the genetic loci and the set of putative core biological processes they affect, and a map explaining how these core processes determine the observed phenotypes. Using simulated data, we demonstrate that SSD can accurately recover the true locus-process and process-phenotype maps as long as at least one of them is sparse. We then apply the method to three empirical datasets, which include the fitness effects of adaptive mutations in different growth conditions, robustness of gene knockouts to a set of genotoxic agents, and the fitness effects of QTLs identified in a yeast cross.

SSD is a flexible method which offers a range of solutions that correspond to different strengths of the sparsity constraints on the locus-process and process-phenotype maps (formally, one unique solution per choice of the hyper-parameters that enforce sparsity). This choice could be made based on some prior biological expectations, or by using standard statistical approaches such as cross-validation to find the set of hyper-parameters that minimizes generalization error. However, since our goal is to identify biologically meaningful low-dimensional structure rather than minimize generalization error, we explore the space of solutions found by SSD across a range of hyper-parameters, and use the reconstruction error landscape and proposed rotation tests to guide the examination of specific solutions. We do not prescribe a method for selecting a single solution. Instead, by exploring solutions with different levels of sparsity, we can examine features of the solutions which are robust to the choice of specific hyper-parameters.

Of course, the use of sparsity as the guiding constraint in our SSD method is a choice, and it would certainly be possible to identify alternative lower-dimensional decompositions of a given dataset by choosing a different set of objectives and constraints. Our choice of sparsity is guided by two main factors. First, because we can use rotation tests to provide evidence for sparsity, we can demonstrate whether or not this constraint is appropriate directly from empirical data (and in cases where there is no evidence for sparsity, SSD should not be used). Second, intuitive notions of modularity in biological systems suggest that sparsity in **M** and **W** may reflect characteristic features of biological organization. For example, sparsity in the locus-process map may reflect a situation where each gene participates in one or a few biological “modules” with specific defined functions, and each such module relies primarily on a relatively small fraction of all possible genes. Sparsity in the process-phenotype map may hold less generally, but could reflect scenarios where any observed phenotype typically depends primarily on a subset of all possible modules. We also note that our method only requires sparsity in one of these two maps, so it could be useful in scenarios where **W** is sparse and **M** is not, or vice versa.

Naturally, even in scenarios where a biological system has a modular structure and sparsity seems intuitively appropriate, all biological processes are inherently coupled at some level. For example, the “omnigenic” model recently introduced by [30] suggests that most loci affect almost every complex trait. The omnigenic model reflects the observation that large numbers of small-effect loci often dominate the heritability of complex traits. This is not inconsistent with the sparsity-inducing *ℓ*_1_ constraint used in SSD. Formally, the *ℓ*_1_ constraint reflects a prior assumption about the distribution (i.e., the spread) of effect sizes, namely, that a small subset of loci have much larger effect sizes than most other loci that affect each process. In contrast, an *ℓ*_2_ constraint, for example, imposes a prior with a tighter spread of effect sizes. This constraint will instead lead to a dense (and non-unique) set of solutions. The sparsity assumption thus remains valid as long as the effects of mutations in the core genes of a pathway are significantly larger than the small effects of the genes outside the pathway, even if there are so many such small-effect genes that they dominate the heritability of the trait.

By using sparsity as a key constraint, our approach produces a different lower-dimensional latent structure in the data than singular value decomposition (SVD), a commonly used method which finds the subspace of a chosen dimensionality that achieves the lowest error in reconstructing the effects matrix (without any additional constraints). By construction, SVD produces a set of processes (formally, basis vectors that span this subspace) which are orthogonal and which are ordered monotonically based on the variation explained by each process. Previous work [16] has shown that SVD applied to a subset of mutations and similar environments generalizes to a held-out set of mutations and dissimilar environments, which suggests that SVD can be fruitfully used to identify an appropriate low-dimensional subspace of processes. However, any set of independent basis vectors which span the subspace will lead to the same generalization error. That is, even though SVD achieves good generalization performance by finding the optimal lower-dimensional decomposition of the genotype-phenotype map, it does not necessarily lead to a unique set of biologically meaningful processes.

Our approach is similar in spirit to Webster, a method based on graph-based dictionary learning introduced recently by Pan *et. al*. [18]. Like SSD, Webster relies on a penalized matrix decomposition framework to identify the locus-process and process-phenotype maps. However, Webster imposes a hard constraint that each locus affects at most two processes and imposes no sparsity constraint on the process-phenotype map. In contrast to Webster, SSD finds sparser solutions with an equivalent reconstruction error, and variable degrees of pleiotropy across loci.

We emphasize that the processes identified by SSD or any other method are fundamentally constrained by the genotypes we study and the phenotypes we choose to measure. We cannot hope to resolve any effects of loci that do not vary across the genotypes we analyze. Thus, it is important to consider the nature of the genetic variation in a given study in interpreting the results of an SSD decomposition: if a given type of variant is not represented, we may fail to identify core processes which depend on those variants. Moreover, it is important to note that expanding a dataset by including additional genotypes can in principle change the inferred structure.

Similarly, the constant effects of loci on all the measured phenotypes are represented by the **b** term in SSD. This reflects the effects of loci on phenotypes that cannot be resolved by the variation in the measured phenotypes. For example, if some core process influences a given type of stress response and we did not measure any phenotypes that depend on that particular type of stress, we would expect the effects of this core process to be absorbed into **b** along with all other processes whose effects do not vary across the measured phenotypes. By measuring additional phenotypes, we could hope to begin to resolve these processes, though our success in doing so would depend on the phenotypes chosen.

We note that by using a matrix decomposition framework, we have implicitly made several important assumptions about the structure of the genotype-phenotype map. First, we have ignored the effects of interactions between loci on the core processes. In other words, we assume that the effect of each locus on each core process does not depend on other loci. Second, the process-phenotype map is assumed to be a linear function of the core processes. Nonlinear structure in the locus-process and process-phenotype maps will lead to structured epistasis between loci in the genotype-phenotype data. This structure is in principle resolvable by measuring epistatic effects between loci for different phenotypes. However, we have focused here on the additive effects matrix, because this is both simpler and can be more reliably estimated given the scope of current data sets.

Finally, our study and others [16, 18] assume a strictly hierarchical genotype to process to phenotype map. That is, we assume that the genotype determines the core processes, which in turn determine the observed phenotypes. This structure has some intuitive appeal, and it is central to any latent structure discovery method of this type. However, it may not always hold in reality. For example, one can imagine a scenario where the effects of mutations on one core process depend on the state of another core process (in other words, core processes affect mutational effects in addition to phenotypes). Our method (along with other matrix decomposition approaches such as SVD) is fundamentally unsuited to describe such scenarios, and developing methods to infer the structure of this and other more general types of genotype-phenotypes maps is an important goal for future work.

## Availability of code and data

Our code, including a tutorial, is available at https://github.com/spetti/sparse-structure-discovery. The three previously published data sets used in this work are accessible from refs. [16, 20, 27].

## Acknowledgements

We thank Andrew Murray and members of the Desai lab for useful discussions. We thank Eliot Fenton for giving us access to his scripts to help process the data from the yeast cross experiments described in Section III C. S.P and G.R were partially supported by the NSF-Simons Center for Mathematical & Statistical Analysis of Biology at Harvard (award number #1764269) and the Harvard Quantitative Biology Initiative. M.M.D acknowledges support from the Simons Foundation (grant 376196), NSF Grant PHY-1914916, and NIH grant R01GM104239.

## SUPPLEMENTARY INFORMATION

### 1. Definitions

1. *Average processes per locus*. Total number of non-zero values in **M** divided by the number of columns in **M** with at least one non-zero value. This definition excludes loci that affect no processes.
2. *Average processes per phenotype*. Total number of non-zero values in **W** divided by the number of rows in **W** with at least one non-zero value. This definition excludes phenotypes that use no processes apart from the linear term **b**.
3. *k-component SVD decomposition*. As with our SSD method, we include a linear term in our decomposition to capture the effects of the loci that do not vary across phenotypes. Given **F**, we let **b** be the mean effect of each locus across phenotypes (i.e. the *L*-dimensional vector where the *i*^*th*^ value is the mean of the *i*^*th*^ column of **F**). Given the SVD of a matrix **F** − **b** = *U* Σ*V* ^*T*^, the *k*-component SVD decomposition of **F** − **b** has **M** equal to the first *k* rows of *V* ^*T*^ and **W** equal to the first *k* columns of *U* with each column scaled by the corresponding diagonal element of Σ. The processs, expressed as *L*-dimensional vectors (rows of **M**), are of unit length, as is the case for decompositions found by our SSD method.
4. *Reconstruction error*. The reconstruction error of the approximation **F** ≈ **WM** + **b** is the squared Frobenius norm of the difference between **F** and the approximation divided by the number of entries in 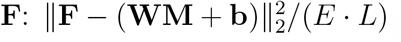.
5. *Cosine error*. To compare the similarity of a decomposition 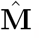, **Ŵ** to the true decom-position **M, W** (for synthetic data), we first adjust 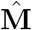 and **Ŵ** to best align the core processes. First, we select the pair of rows **M**_*k*,:_ and 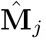 with the highest absolute value of cosine between them and assign 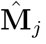 and **Ŵ**_:,*j*_ to the *k*^*th*^ row and column of the adjusted matrices 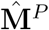 and **Ŵ**^*P*^ respectively. Further if the cosine between **M**_*k*,:_ and 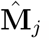 is negative, we multiply the *k*^*th*^ row and column of 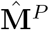 and **Ŵ** ^*P*^ (respectively) by negative one. We repeat this process, excluding the rows in **M** and 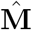 that have already been paired. This process permutates and changes the sign of the core processes, but does not change the approximation: 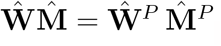.

The mean cosine error for **M** measures the similarity between the pairs of corresponding core processes, viewed as *L*-dimensional vectors: 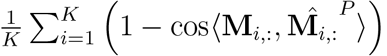.

The mean cosine error for **W** measures the extent to which each phenotype uses the corresponding processs similarly: 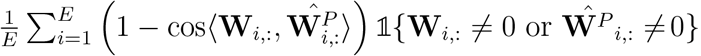. The indicator function ensures that the phenotypes affected by no core processes (other than the linear term) in both the true and predicted decompositions do not contribute to the error.

### 2. Sparse Structure Discovery

SSD takes as input the additive effects matrix **F**, an upper bound on the desired number of processes *K*_max_, and the regularization parameters *λ*_*W*_, *λ*_*M*_. It returns **M, W** and **b** that approximately minimize

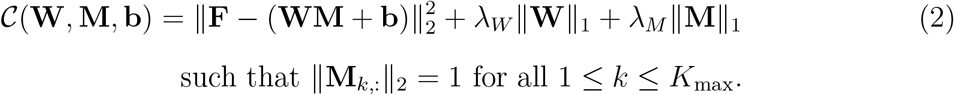

Initially **b** is set to the column means of **F**, and **W** and **M** are found by taking the Singular Value Decomposition (SVD) of **F**−**b** with the top *K*_max_ singular vectors. We then alternately fix **W** and find **M** and **b** that optimize (2), (ii) normalize the rows of **M**, (iii) fix **M** and **b** and find **W** that optimizes (2). While the objective function (1) is not jointly convex in **W** and **M**, the optimization problems in (i) and (iii) are each convex and can be efficiently solved.

In order to use comparable regularization values and obtain comparable errors across input matrices **F** with different sizes and magnitudes, we normalize the input matrix **F** before performing SSD and the rotation tests. To normalize **F**, we divide each entry by the standard deviation of all the values in **F**. In both the SSD solution space plots and the rotation tests, the reported reconstruction error is with respect to this normalized version of **F**. After normalization, the reconstruction error can be interpreted as the fraction of variance unexplained by the decomposition.

For each application, we apply our method with 625 pairs of regularization parameters: 25 values of *λ*_*W*_ uniformly distributed between 10^−3^ and 1.5 in logscale and 25 values of *λ*_*M*_ uniformly distributed between 10^−4^ and 10^−2^ in logscale. We choose *K*_max_ as the minimum number of SVD components needed to explain at least 95% of the variance in **F** − **b**. Choosing *K*_max_ *<* min{*E, L*} speeds up the method. Recall that the optimization procedure automatically picks an appropriate number of processes *K* ≤ *K*_max_ for a given *λ*_*M*_, *λ*_*W*_.

#### a. Comparison to other penalized matrix decomposition methods

Our Sparse Structure Discovery method is a form of penalized matrix decomposition. It is well-known that the low rank matrix decomposition that gives the best approximation of a matrix with respect to the Frobenius norm can be computed via the singular value decomposition (SVD) (see [31]). Penalized matrix decomposition refers to a broader range of matrix decomposition formulations whose objectives are to both minimize the Frobenius norm of the approximation and to encourage the matrix factors to exhibit particular properties (e.g. sparsity) through hard constraints or regularization [7].

One form of penalized matrix decomposition is *sparse coding*, where the goal is to identify an overcomplete set of basis vectors, often called dictionary elements, so that each data point can be written as a combination of a small number of dictionary elements. This approach was used by Field and Olshausen to identify putative receptive fields of cells in the visual cortex [9]. The computer science and statistics literature has developed various formulations of sparse coding and accompanying efficient algorithms for finding the dictionary elements [8]. Algorithms for sparse coding formulations that impose an *L*_0_ penalty on the use of dictionary elements are studied in [19, 32, 33]. Algorithms for the more tractable convex relaxation with an *L*_1_ penalty are studied in [34–36]. In Appendix 6 we further discuss the graph-regularized approach introduced in [19] and applied to the genotoxin data set in [18].

The key difference between SSD and sparse coding is that we enforce sparsity in both the dictionary elements (**M** matrix) and the description of the data as combinations of the dictionary elements (**W** matrix). This is motivated by our observation that sparse solutions can be found for both **W** and **M** in empirical genotype-phenotype maps with a marginal increase in reconstruction error. In contrast, standard sparse coding approaches do not constrain the sparsity of the dictionary elements. Additionally, the vector **b** in (2) is introduced to capture the effects of loci on processes that do not have a variable effect on the measured phenotypes.

### 3. Rotation tests for locus and phenotype sparsity

For both tests, we first subtract out the mean effect of each locus across phenotypes to approximate **b**, the effects that do not vary across phenotypes. Then, we normalize **F** and select *K*_max_ as described in Section 2. For the locus-rotation test, we rotate the rows of **F** randomly by right-multiplying by a random *L*×*L* orthogonal matrix **O** drawn from the Haar distribution, as implemented by SciPy’s stat.orthogroup library. We apply our SSD method directly to **FO** (without normalizing) for 25 values of *λ*_*M*_ uniformly distributed between 10^−4^ and 10^−2^ in logscale and *λ*_*W*_ = 10^−3^. For the phenotype-rotation test, we left-multiply **F** by a random *E* × *E* orthogonal matrix **O**^′^ drawn from the Haar distribution and apply SSD to **O**^′^**F** for 25 values of *λ*_*W*_ uniformly distributed between 10^−3^ and 1.5 in logscale and *λ*_*M*_ = 10^−4^.

### 4. Synthetic data with hub-and-spoke structure

To test our SSD method on data with more complex underlying structure, we generated synthetic data with a hub-and-spoke structure, as illustrated in Figure S2a. We constructed eight *H*-processes and four *P* -processes. Each of *L* = 200 loci participated in each process independently with probability 0.2, the weights of the participating loci were drawn independently from a standard normal, and the rows of **M** were normalized. We then constructed 20 groups of 5 phenotypes: one hub phenotype and four perturbations of the hub phenotype, which we call spokes. The hub phenotype depends on two randomly selected *H*-processes with weights drawn independently from a standard normal. Each spoke phenotype is a sum of the hub phenotype and one of the four *P* -processes multiplied by a scaling factor drawn from a standard normal. This construction yields a 100 ×12 matrix **W**, a 12 ×200 matrix **M** and the fitness effect matrix **F** = **WM** + *η*, where the noise *η* is drawn independently from a normal distribution with scale 0.3 times the standard deviation of the entries in **WM**.

This same **F** can also be expressed as a decomposition 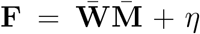 with 24 core processes and with more sparsity in 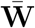 than **W** and far less sparsity in 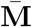 than **M**, see Figure S2b. To obtain 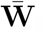, we keep the four *P* -processes and construct an *S*-process for each of the 20 hub phenotypes. Instead of expressing each hub phenotype as the weighted sum of two *H*-processes, each hub phenotype is now represented by single *S*-process.

### 5. Analysis of adaptive mutations in yeast (Kinsler et. al. dataset)

The dataset in Kinsler et. al. [16] contains the additive effects of 421 adaptive mutations in 45 environments. We chose a subset of 288 mutations using the procedure described in the original work. Specifically, mutations that were either not sequenced, whose mean additive effect across the 8 evolutionary conditions was smaller than a threshold (0.05) or whose maximal error of the additive effect over all environments was larger than a threshold (0.5) were removed. The specific thresholds were not specified in Kinsler et. al.; we chose thresholds such that we were left with close to the total number of mutations analyzed in this work (i.e., 292).

#### a. Clustering mutations

Clustering of the **M** matrix was performed through hierarchical/agglomerative clustering [37] (using the linkage function in SciPy’s hierarchical clustering library) with an absolute cosine metric. Since our goal was to cluster loci with similar effect profiles on processes (i.e., columns of **M**) independent of the overall sign and magnitude, we use a metric 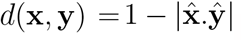, where **x, y** are two vectors and 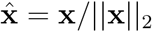 denotes the unit vector. The method groups the loci into clusters depending on an input distance threshold. We found that for a large range of thresholds (0.15 to 0.93), the number of clusters ranged from 6 to 11. There was no sharp delineation within this range. We chose an intermediate threshold value 0.4, which led to 8 clusters. For the analysis with fewer diploids (Figure S4) we used a threshold value of 0.22 to obtain 8 clusters.

In Figure S5d, we present results from hierarchical/agglomerative clustering of the **M** found using SVD. We chose a distance threshold of 0.47 instead of 0.4 for the SSD solution in Figure 3d,e to obtain 9 clusters since we could not find a threshold which led to 8 clusters. Choosing a matching threshold of 0.4 led to 11 clusters.

#### b. Bi-cross-validation

In this Section, we summarize the bi-cross-validation test described in [38] and applied in [16]. We split the 45 environments into train and test environments, and the 288 mutations into train and test mutations. In panel a of Figure S5, the train and test environments are the subtle (25) and strong perturbation (20) environments as defined in [16], respectively (Recall that environments in which the fitness effects differed slightly and significantly from the average fitness effects in the evolution condition were classified as subtle and strong perturbations respectively). In panel b, the training and test environments are chosen randomly in a 36:9 split.

Each result is averaged over eight random splits of the mutations into training and test sets. In each random split, the training set contains 60 training mutations and test set contains 228 test mutations. To split mutations, the number of mutations of each annotation (Diploids, IRA1-mis, IRA2, etc) that are included the training and test sets are decided as specified in [16]. The specific mutations assigned to each set are sampled randomly. For example, Kinsler et. al. assign 20 diploids to the training set and 168 diploids to the test set. The specific set of 20 diploids that are assigned to the training set for each of the 8 random seeds are sampled with equal probability from the full set of 188 diploids. As described in [16], the weighted reconstruction error is computed by normalizing the total reconstruction error for all mutations of an annotated class with the number of mutations in that class. This ensures that the performance on the diploids are not overrepresented in the results.

To obtain the bi-cross validation reconstruction error for each method, we first decompose **F** on train environments and mutations into two matrices **W**_1_, **M**_1_. Fixing **M**_1_, we fit the process-phenotype map **W**_2_ for the test environments and train mutations. Similarly, fixing **W**_1_, we fit the locus-process map **M**_2_ for the test mutations and train environments. The predicted loci-phenotype map on test environments and mutations is then **W**_2_**M**_2_. To compare SVD and SSD on an equal footing, we first subtract from **F** the mean of **F** across environments for each locus.

### 6. Comparison to Webster method on genotoxin dataset

SSD differs from Webster in three key ways. First, Webster imposes locus-sparsity as a hard constraint; each locus particpates in at most *j* core processes where *j* is an input parameter. In contrast, SSD allows loci to participate in different numbers of core processes, allowing the loci to exhibit varying degrees of pleiotropy. Second, whereas phenotype-sparsity is a tunable parameter in SSD, Webster does not enforce phenotype-sparsity. Finally, Webster’s optimization includes graph regularization objectives that encourage each locus to have a similar core process membership profile as its five closest neighbors, and analogously for phenotypes. This arbitrary cutoff of five could cause problems for a locus or phenotype that is significantly dissimilar from all others.

We first select SSD solutions to compare to the Webster decomposition of the genotoxin dataset [20] presented in [18]. In the Webster decomposition each locus participates in exactly two of ten core processes. We selected the most comparable SSD solution (ten processes, 2.0 average-processes-per-loci, 6.8 average-processes-per-genotoxin), as well as the SSD solution we selected using the rotation tests as a guide (illustrated by the white star in Figure 4, ten processes, 3.3 average-processes-per-loci, 2.5 average-processes-per-genotoxin), and the 10-process SVD solution.

Next, we compared the unnormalized reconstruction error for each genotoxin between the two SSD solutions described above, the SVD decomposition, and the Webster method (left column in Figure S6). Predictably, the methods with less strict sparsity requirements give lower mean error (SVD 1.4, selected SSD solution 2.6, most comparable SSD solution 2.9, Webster 3.3). Unlike Webster, our SSD method allows the number of processes that each locus participates in to vary, reflecting the possibility that loci may exhibit different levels of pleiotropy (right column Figure S6). This flexibility may account for the improved predictions of our SSD solutions over Webster at the same average locus-sparsity. As displayed in Figure S6 center column, the process-genotoxin maps from the SSD solutions are more sparse.

Of the sparse solutions, our selected SSD solution most accurately reconstructs the additive effects matrix and exhibits the most genotoxin-sparsity (see Figure S6, center column). Moreover, the locus-sparsity of this solution is sufficient to assign putative biological functions for many of the core processes using predefined annotations (Figure 4d). This suggests that the SSD approach is a more promising method for generating biologically reasonable hypotheses about genetic architecture in this system.

### 7. Joint QTL mapping from large-scale genotype-phenotype measurements

Given an *E*×*N* matrix **Y** encoding *E* measured phenotypes of *N* individuals and an *S*×*N* {0, 1}-valued matrix **X** expressing the genotypes of the *N* individuals at *S* loci, our joint QTL mapping method identifies *L < S* putative causal loci which explain the majority of the predictable variation in the measured phenotypes. The output of our method is an *E* ×*L* effects matrix **F** which approximates the phenotypes as an additive function of the effects of these *L* loci. The key step in our method aligns loci across phenotypes using a penalized regression framework based on *ℓ*_2,1_ regularization with a highly optimized implementation called *glmnet* [39]. Specifically, we minimize

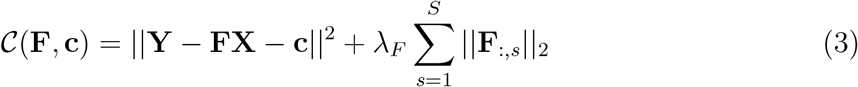

with respect to **F** and **c**, where 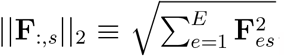, *λ*_*F*_ controls the strength of regularization and **c** is an *E* × 1 intercept term. This *ℓ*_2,1_ regularization penalty is a generalization of the well-known *ℓ*_1_-based Lasso to multiple outcomes. Like Lasso, the *ℓ*_2,1_ penalty favors sparse solutions by selecting only the loci whose effects across phenotypes (as measured by ||*F*_:,*s*_||_2_) are sufficiently large, thus automatically identifying and aligning both large-effect, non-pleiotropic loci and loci that have small effects across many phenotypes.

In our yeast cross application, we have *N* ≈ 100, 000 segregants and *S* ≈ 42, 000 loci. Due to the scale of this data and strong correlations between neighboring loci from linkage, we avoid running *glmnet* on all 42, 000 loci. We instead run *glmnet* on a smaller subset of putative causal loci and develop a statistical method for computing confidence intervals to narrow down the true locations of each causal locus. Our pipeline is as follows:

1. Compute a reduced genotype matrix by restricting **X** to a set of rows corresponding to loci that are pairwise correlated by no more than 94%. On our yeast dataset, this reduces *S* from ≈ 42, 000 to 1579.
2. Perform *ℓ*_2,1_-regression on the reduced genotype matrix. On our yeast dataset, this yields 1314 putative causal loci (non-zero columns of **F**).
3. Construct a new list of putative casual loci that are more likely to be casual than the loci selected in Step 1. To do so, compute confidence intervals for each putative casual locus for each phenotype separately using the statistical method described in Section 7 a. When the confidence intervals for a single locus do not overlap, it suggests that the locus is summarizing the effect of multiple distinct nearby causal loci. We “split” the locus by adding a set of loci to the new list such that each phenotype’s confidence interval contains at least one locus in the set. When the confidence intervals for a locus overlap across all phenotypes, we add the locus from the intersection with the strongest evidence of being causal to the new list. The same locus may appear multiple times on the new list, suggesting that the *ℓ*_2,1_ optimization assigned the effect of a single locus to two (or more) nearby loci. After removing such redundancies, the new list contains 1119 loci for our yeast dataset.
4. Perform *ℓ*_2,1_-regression on the genotype matrix restricted to the new list of putative casual loci. We use this **F** in downstream analysis. On our yeast dataset, this yields 1089 putative casual loci (non-zero columns of **F**).
5. Localize the ORFs of the putative causal loci with the strongest effects by computing confidence intervals for each phenotype.

In Step 1, we apply a greedy algorithm to pre-filter the loci. We order the SNPs (loci) by genomic position. We select the first SNP. We subsequently select the next SNP that has genotypic (Pearson) correlation *<* 0.94 with the most recently selected SNP. This process is repeated until we get to the last SNP.

Steps 2 and 4 use the implementation of *ℓ*_2,1_-based regression from the *glmnet* R library [39]. The regularization parameter *λ*_*F*_ in Eq. (3) is set using cross-validation. Specifically, the training, validation, and test sets are obtained by splitting the columns of **X** (corresponding to segregants) in the ratio 80:10:10, *glmnet* solves Eq. 3 on the training set for a range of *λ*_*F*_, and we select the solution with the minimum mean absolute error on the validation set. We use the test set to evaluation our predictions before and after matrix decomposition, see Figure S7a.

The goals of Step 3 are to more accurately localize the putative causal loci returned by Step 2 and to determine whether some putative causal loci are summarizing the effects of multiple nearby loci with distinct effects. The putative causal loci identified by *glmnet* in Step 2 are a subset of the loci chosen via the greedy prefiltering done in Step 1. Therefore, it is quite possible that the true causal locus was filtered out in Step 1, and the putative causal locus identified by *glmnet* is a nearby locus that is highly correlated with the true casual locus. Alternatively, a putative causal locus identified by *glmnet* may describe the effect of one nearby causal locus for certain phenotypes and a different nearby causal locus for other phenotypes, i.e. the putative causal locus is summarizing multiple loci with different effects.

To arrive at a new list of loci that we believe to more likely to be causal, we replace each locus identified in Step 2 with a set of loci constructed according to the following procedure. For each locus *ℓ*, we first apply the method described in Section 7 a to compute a confidence interval of locations for the true causal locus for each phenotype separately. For each locus *z* in the confidence interval for phenotype *e*, we also return best approximation of the linear effect of locus *z* on phenotype *e*, which we denote 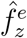 (computed as described above Eq. 7). Across loci in a confidence interval, a higher value of 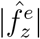 indicates that locus *z* is more likely to be causal.

We iteratively select loci for the new set as follows. For each locus *z* in some confidence interval, we compute 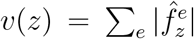 where 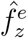 is set to zero when locus *z* is not in the confidence interval for phenotype *e*. The locus *z*^*^ that maximizes *v* is added to the new set. If locus *z*^*^ is in the confidence interval for all phenotypes, we add no other loci to the new set. Effectively, we have replaced *ℓ* with a nearby locus *z*^*^ that is in the confidence interval for each phenotype and exhibits a stronger effect (measured by the magnitude of effect size summed across environments). If there are phenotypes whose confidence intervals do not contain *z*^*^, it is likely the case that locus *ℓ* summarizes the effects of different causal loci for different phenotypes. We need to include more loci in the new set so that the new set includes a least one locus in the confidence interval of each phenotype. To do so, we remove all phenotypes whose confidence intervals contain *z*^*^ and again find the locus *z*^**^ that maximizes *v* (where now the summation in *v* is over a restricted set of phenotypes whose confidence intervals do not contain *z*^*^). We repeat this process until the each confidence interval contains at least one locus in the new set.

In Step 5 we localize the putative causal loci returned by *glmnet* in Step 4 to ORFs. Pinpointing the location is only possible for the strongest effect loci, so we restrict our analysis to loci that exhibit an additive effect of magnitude at least 0.003 for some phenotype.

For each such locus *ℓ* and phenotype *e*, we again use the method described in Section 7 a to compute a confidence interval and the best approximations of the linear effects 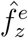 for each locus *z* in the confidence interval. For each locus *z* in some confidence interval, we again compute 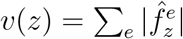 where 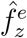 is set to zero when locus *z* is not in the confidence interval for phenotype *e*. We declare the locus *z*^*^ that maximizes *v* the “top” locus. We consider the intersection of all confidence intervals containing *z*^*^ to be the common confidence interval for locus *ℓ*. We label locus *ℓ* with the names of all ORFs corresponding to a locus in this common confidence interval.

#### a. Confidence interval computation

We describe a method to identify a confidence interval for a single locus with respect to a single phenotype. We assume a linear model for the effect of a locus on the phenotype of segregant *n*

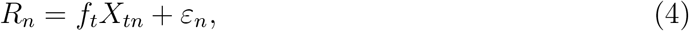

where *R*_*n*_ is the “residual”, i.e., the phenotype measurement not explained by the rest of the loci, *t* is the index of the true locus, *f*_*t*_ is its true fitness effect, and *ε*_*n*_ ∼ 𝒩(0, *σ*^2^) is a noise term which is drawn i.i.d from a normal distribution with mean zero and variance *σ*^2^. To measure how well a nearby locus *z* explains the residuals, we compute the squared error between the observed residuals and the best approximation of the residuals as a linear function of *X*_*z*,:_, which we call 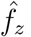. We define this error as

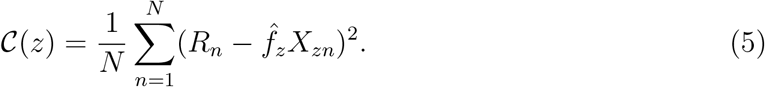

To arrive at a confidence interval, we suppose that *ℓ* is the locus that minimizes 𝒞 when *t* is the true causal locus and compute the probability that *ℓ* minimizes *C* under this assumption:

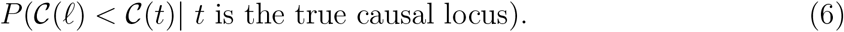

If this probability is less than 0.023 (two standard deviations), we reject the hypothesis that *t* is the true causal loci and exclude *t* from the confidence interval.

Now we explain how to compute (6). Let 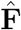 and **ĉ** be the putative additive effects matrix and linear term returned by *ℓ*_2,1_ optimization, and let *ℓ* and *e* be the locus and phenotype of interest respectively. Since we consider one phenotype at a time, we suppress the dependency on *e* and write **Y** = **Y**_*e*,:_ and *c* = **c**_*e*_. By a slight abuse of notation, when *ℓ* appears as a subscript of 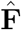 it refers to the column corresponding to the locus *ℓ* and when *ℓ* appears as a subscript of **X** it refers to the row corresponding to the locus *ℓ* (these will not necessarily have the same index). Throughout, we use bar to denote averages over the *N* segregants. We use the putative additive effects map to compute the residuals,

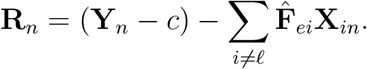

For a locus *z*, the best approximation of the residuals as a linear function of **X**_*z*,:_, i.e. the value of 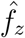 that minimizes 𝒞 (*z*), is 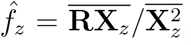. Plugging this expression into Eq. 5, we have 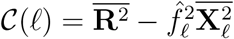 and 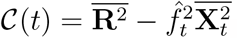. Taking the difference, we obtain

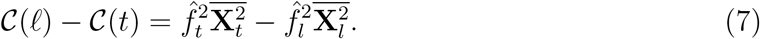

Since **R**_*n*_ = *f*_*t*_**X**_*tn*_ + *ε*_*n*_, it follows that

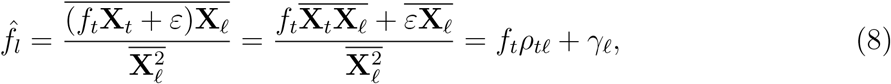

where 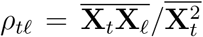 is the fraction of segregants with genotype +1 at *t* that also have genotype +1 at *ℓ*, and 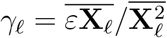 is a random variable equal to the average noise over all segregants with genotype +1 at *ℓ*. Similarly,

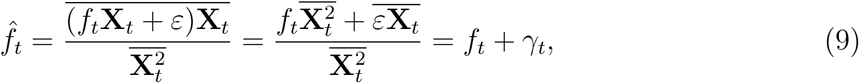

where 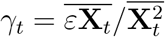 is a random variable equal to the average noise over all segregants with genotype +1 at *t*. Plugging these into Eq. (7), we obtain

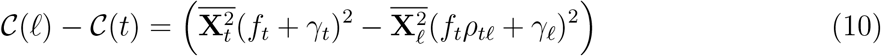

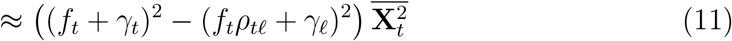

Assuming that *ℓ* and *t* are nearby, linkage guarantees that most segregants will have the same genotype at these positions. As a result, 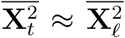 (validating approximation (11)) and *ρ*_*tℓ*_ will be close to one. Since *γ*_*t*_, *γ*_*ℓ*_ are order 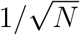 (as they are the mean of order *N* normals with constant variance *σ*^2^), they tend to be much smaller than *f*_*t*_ whenever *f*_*t*_ is significant enough to be causal. We therefore may assume that *f*_*t*_ and *f*_*t*_ + *γ*_*t*_, and *f*_*t*_*ρ*_*tℓ*_ + *γ*_*ℓ*_ have the same sign.

Suppose *f*_*t*_ *>* 0. Then 𝒞 (*ℓ*) *<* 𝒞 (*t*) whenever *f*_*t*_(1 − *ρ*_*tℓ*_) *< γ*_*t*_ − *γ*_*ℓ*_. Let Γ be the random variable equal to *γ*_*t*_ − *γ*_*ℓ*_. Again using the approximation that 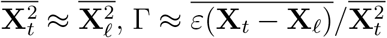 is 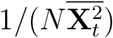 times the difference between the noise summed over all segregants with genotype +1 at *t* and 0 at *ℓ* and the noise summed over all segregants with genotype 0 at *t* and +1 at *ℓ*. (Note Γ is not affected by the noise from segregants that have the same genotype value at *t* and *ℓ*.) Thus, we can approximate Γ as 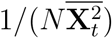 times the sum of *d* i.i.d. draws of 𝒩 (0, *σ*^2^) where 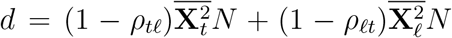 is the number of segregants with a recombination breakpoint between *t* and *ℓ*. The assumption that 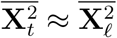 implies *ρ*_*tℓ*_ ≈ *ρ*_*ℓt*_ and 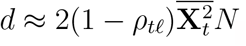. We approximate 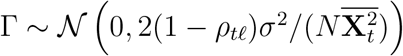. It follows that

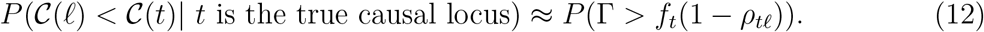

The probability of this event is less than 2.3% whenever the value *f*_*t*_(1 − *ρ*_*tℓ*_) is at least 2 standard deviations of Γ. Thus, we reject the null hypothesis that *t* is the true causal locus whenever

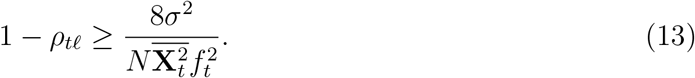

Note we are most likely to reject the null hypothesis when *f*_*t*_ is high (the true locus has a large effect) and *ρ*_*tℓ*_ is small (a high fraction segregants have a breakpoint between *t* and *ℓ*). In other words, it is easiest to identify the causal locus when its effect size is large and there are many segregants with breakpoints nearby.

In practice, to verify (13) we approximate 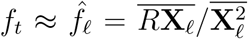 and *σ*^2^ as the cost 𝒞 (*ℓ*). First, as derived in (8), the estimated effect size 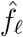 will differ from the true effect size *f*_*t*_ by 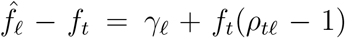. The relative error of the former approximation is 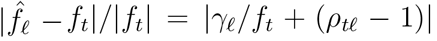, which is small since |*γ*_*ℓ*_| ≪ |*f*_*t*_| and 1 − *ρ*_*tℓ*_ ≪ 1. Second, we have 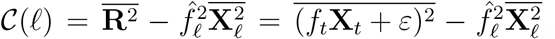. Expanding this expression and using 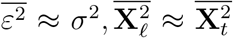 gives 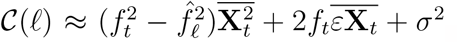. Note that 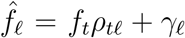 and 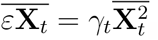. Thus, 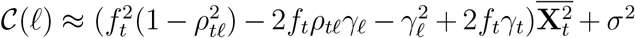. Since 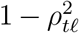 is small and *γ*_*t*_, *γ*_*ℓ*_ are both order 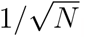, 𝒞 (*ℓ*) is a good estimator for *σ*^2^.

#### b. Comparison to other QTL mapping approaches

Existing approaches for mapping QTLs of multiple traits include composite interval mapping [40], least squares regression [41], and Bayesian inference [42, 43]. See survey given in Chapters 14 and 15 of [44]. The scale of our dataset (∼ 42000 loci, ∼ 100, 000 individuals) renders such methods intractable. Instead, we turn to *glmnet*, a fast solver for regularized generalized linear models [39] that is capable of handling the scale of our data. In [45], Qian et. al. apply *glmnet* with a standard lasso penalty for QTL mapping of four traits separately using data from the UK biobank. We extend this approach by mapping QTLs for multiple traits simultaneously using *glmnet* with an *ℓ*_2,1_ error. Moreover, the extreme linkage present in our dataset necessitates post-processing to identify confidence intervals for the casual loci.

**FIG. S1:**
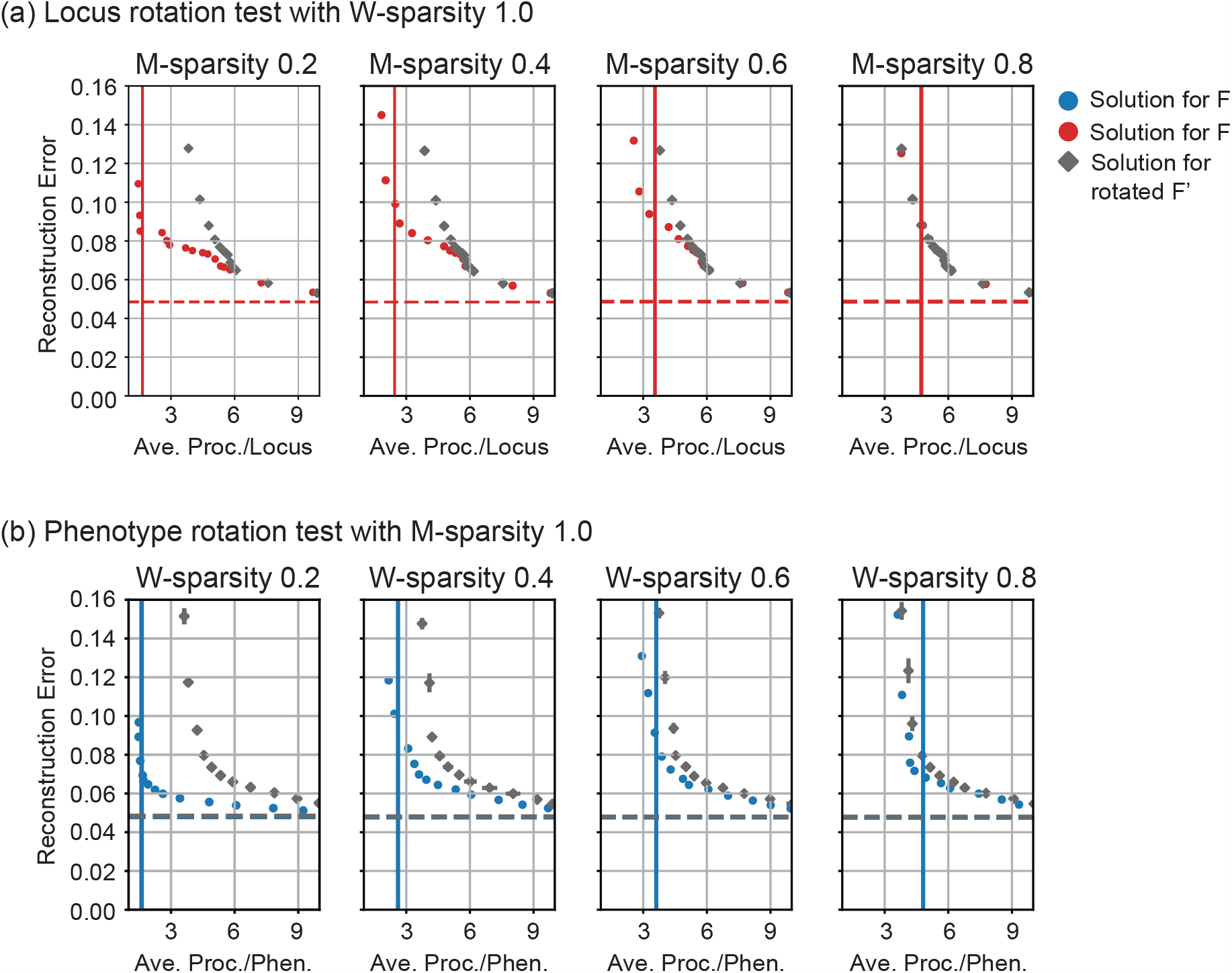
Rotation tests on synthetic data over a range of sparsities. (a) Analogous plots to the loci rotation test in Column 1 of Figure 2 for a synthetic additive effects matrix with a range of **M**-sparsities and **W**-sparsity equal to 1. (b) Analogous plots to the phenotype rotation test in Column 1 of Figure 2 for a synthetic additive effects matrix with a range of **W**-sparsities and **M**-sparsity equal to 1.

**FIG. S2:**
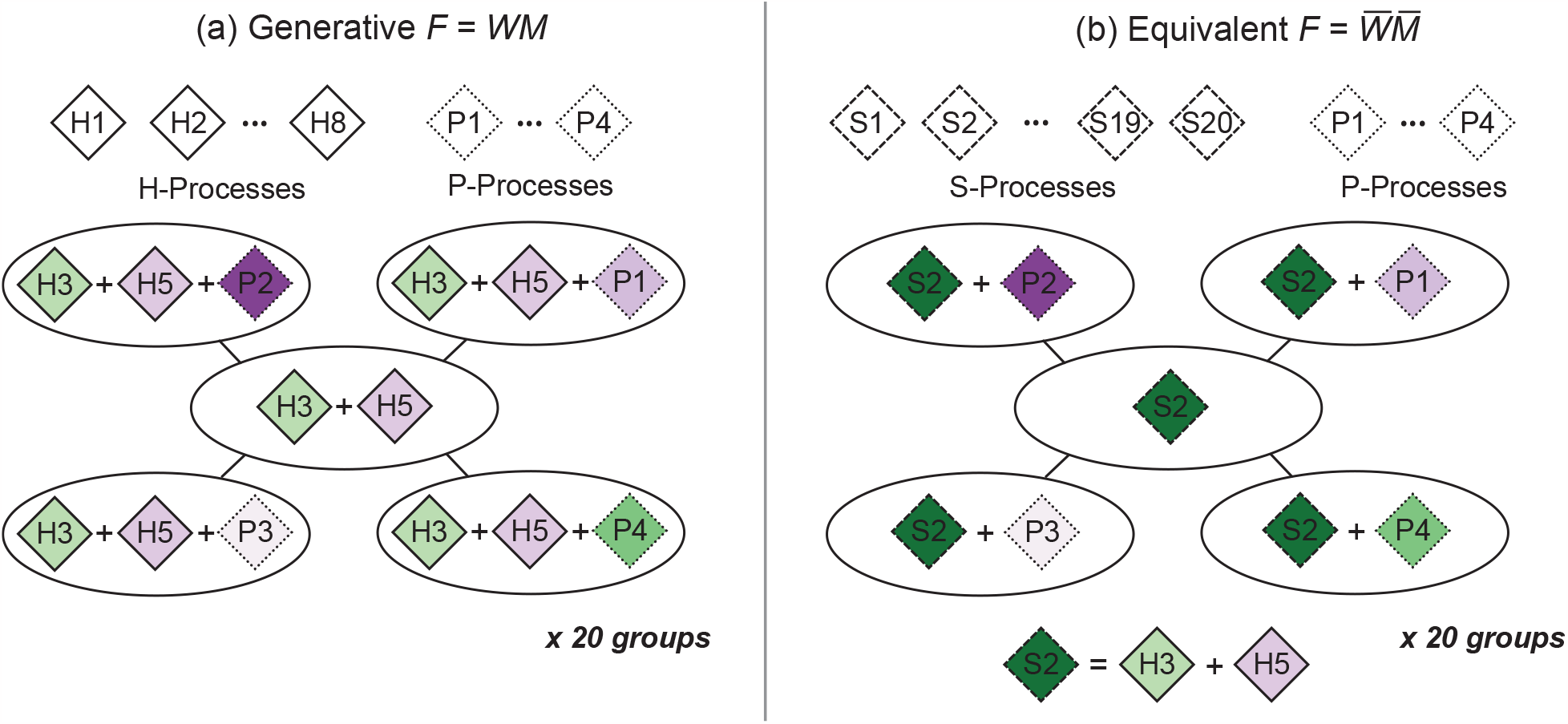
Generation of hub-and-spoke synthetic data. Diamonds represent processes, ovals represent phenotypes, and the color of the process represents its weight in the phenotype. (a) The core phenotype (center hub) is the weighted sum of two hub processes (H-process), and each perturbation is the sum of the processes of the core phenotype plus a weighted perturbation process (P-process). The group of five phenotypes depicted here corresponds to the group of phenotypes labeled in Figure S3c. We generate 20 such groups from the common set of 8 hub and 4 perturbation process, as detailed in Methods 4. (b) An alternate way to generate the same **F** matrix is to replace the *H*-process with one *S*-process per phenotype.

**FIG. S3:**
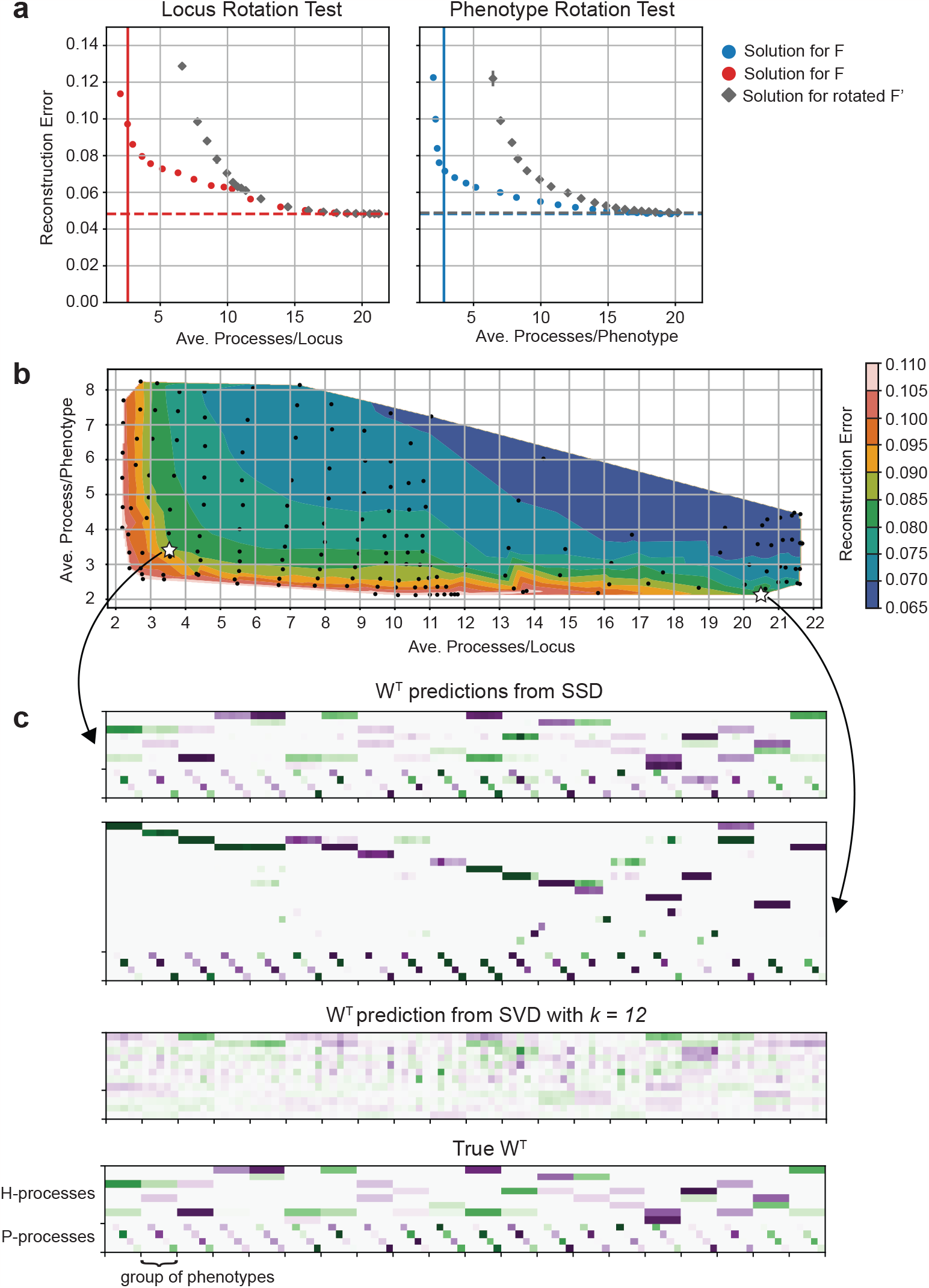
SSD on hub-and-spoke synthetic data. (a) The rotation tests suggest both locus and phenotype sparsity. (b) Our SSD method finds a range of solutions at different sparsity and error levels. We consider two SSD solutions: one with 12 core processes and reconstruction error 0.086 that is sparse in both loci and phenotypes (lower left star) and one with 21 processs and reconstruction error 0.083 that is sparse in phenotype only (lower right star). (c) Illustrations of predicted and true *W*^*T*^. The values are illustrated on a purple-to-green scale ranging from -10 times to +10 times the average magnitude of an entry in the **W** matrix. The five phenotypes labeled “group of phenotypes” are illustrated in Figure S2a. The matrix **W** for the 12 core process solution approximates the generative **W** well. The matrix **W** for the 21 core process solution has a structure similar to alternate generative structure 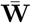 described in Figure S2b.

**FIG. S4:**
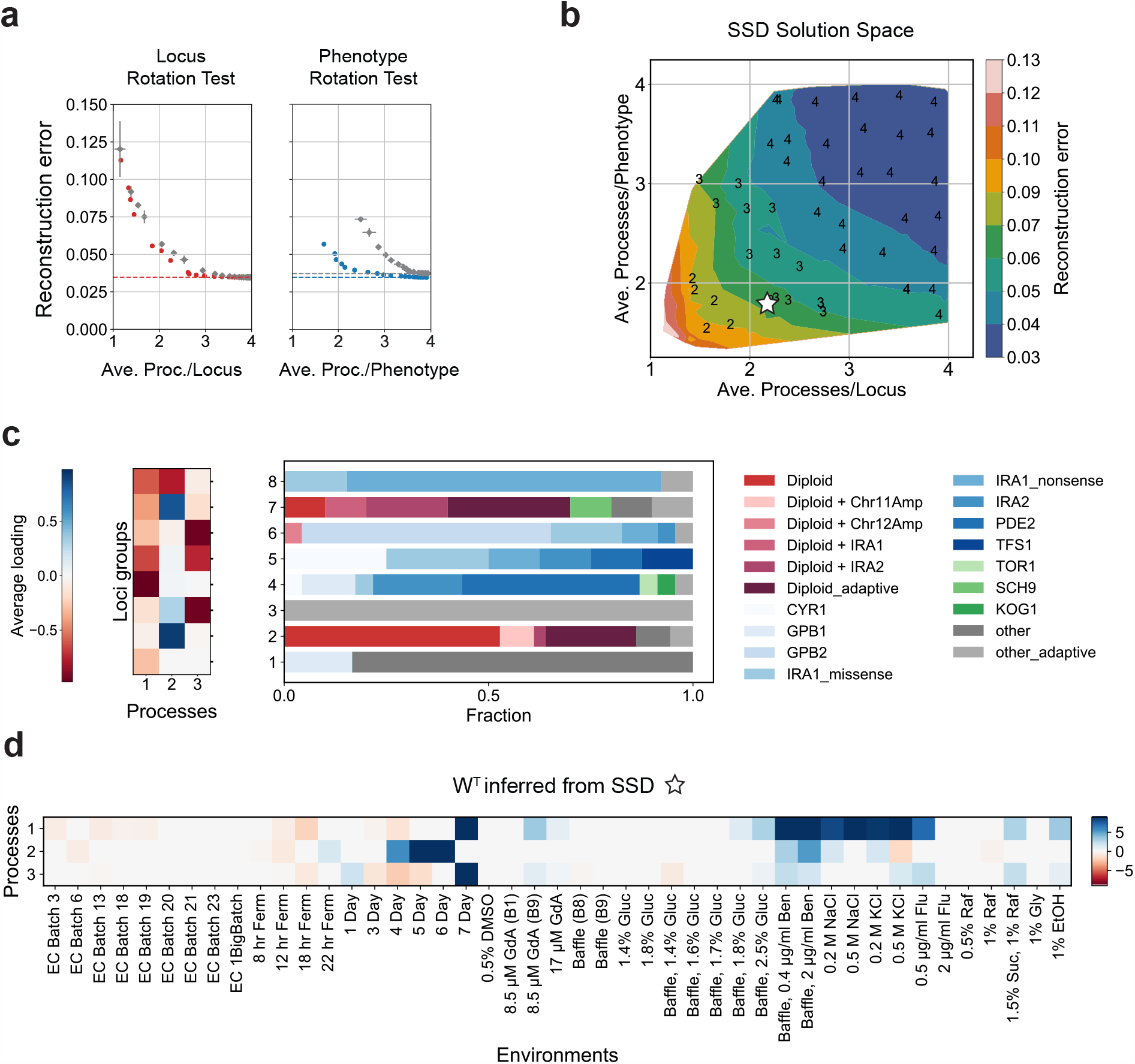
SSD applied to Kinsler et. al. [16] data with fewer diploid mutants. To ensure that the many diploids do not bias our results, we repeated the analysis presented in Figure 3 with a reduced effects matrix **F**. Specifically, we randomly sampled 20 diploids of the 188 in the original dataset leading to an **F** with dimensions 45 × 120. Despite much lower locus-sparsity, the examined **M** and **W** solutions show similar features as the ones obtained using the full **F** (Figure 3). (a) The locus rotation test shows much reduced sparsity in the locus-process map compared to the dataset with diploids included (Figure 3b). The sparsity in the process-phenotype map is retained. (b) The solution space illustrating highly sparse solutions with low reconstruction error. The selected solution (*K* = 3), which is chosen to match the reconstruction error of the solution picked in Figure 3, is marked with a white star. (c) The **M** matrix with loci clustered into 8 groups based on linkage clustering of loci with a modified cosine similarity metric as in Figure 3d. (d) The process-phenotype map **W**. Processes 1 and 2 from the full **F** (Figure 3e) are comparable to processes 1 and 2 respectively, whereas process 3 here appears to capture processes 3 and 4 for the full **F**.

**FIG. S5:**
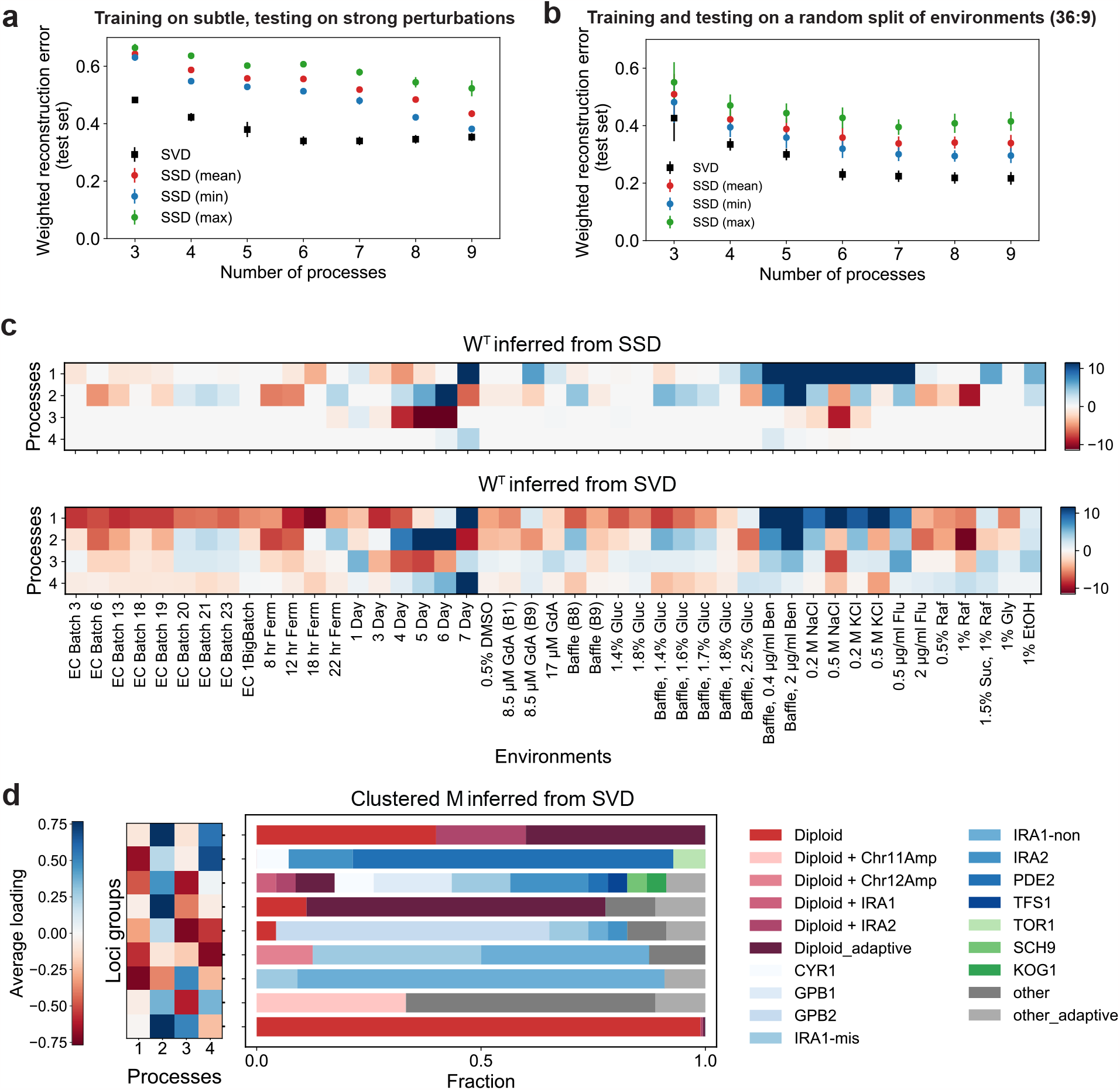
Comparison of SSD and SVD decompositions on Kinsler et. al. [16] data. (a,b) Bi-cross-validation on held-out sets as described in [38] and applied in [16]. See Section 5 for more details. The results are averaged over 8 random seeds. For SSD, we present the minimum, maximum and mean weighted reconstruction errors across all the hyper-parameters *λ*_*W*_, *λ*_*M*_ described in the Methods for a given number of processes *K*. SVD of the same rank tends to show lower generalization error compared to SSD. (c) The process-phenotype map **W** from SSD and SVD, highlighting that the SSD solution is much sparser. The SSD solution is reproduced from Figure 3e. Since SVD does not fit **b** separately, here we estimate **b** as the mean effect across environments for each locus and subtract it from **F** before applying SVD. (In Kinsler et. al., they do not subtract the means, and so their first SVD component approximately represents the constant effect **b**.) (d) Hierarchical/agglomerative clustering of **M** inferred from SVD similar to Figure 3d (see Methods for clustering parameters). Note the denser loading matrix as compared to the analogous figure for the SSD solution (Figure 3d).

**FIG. S6:**
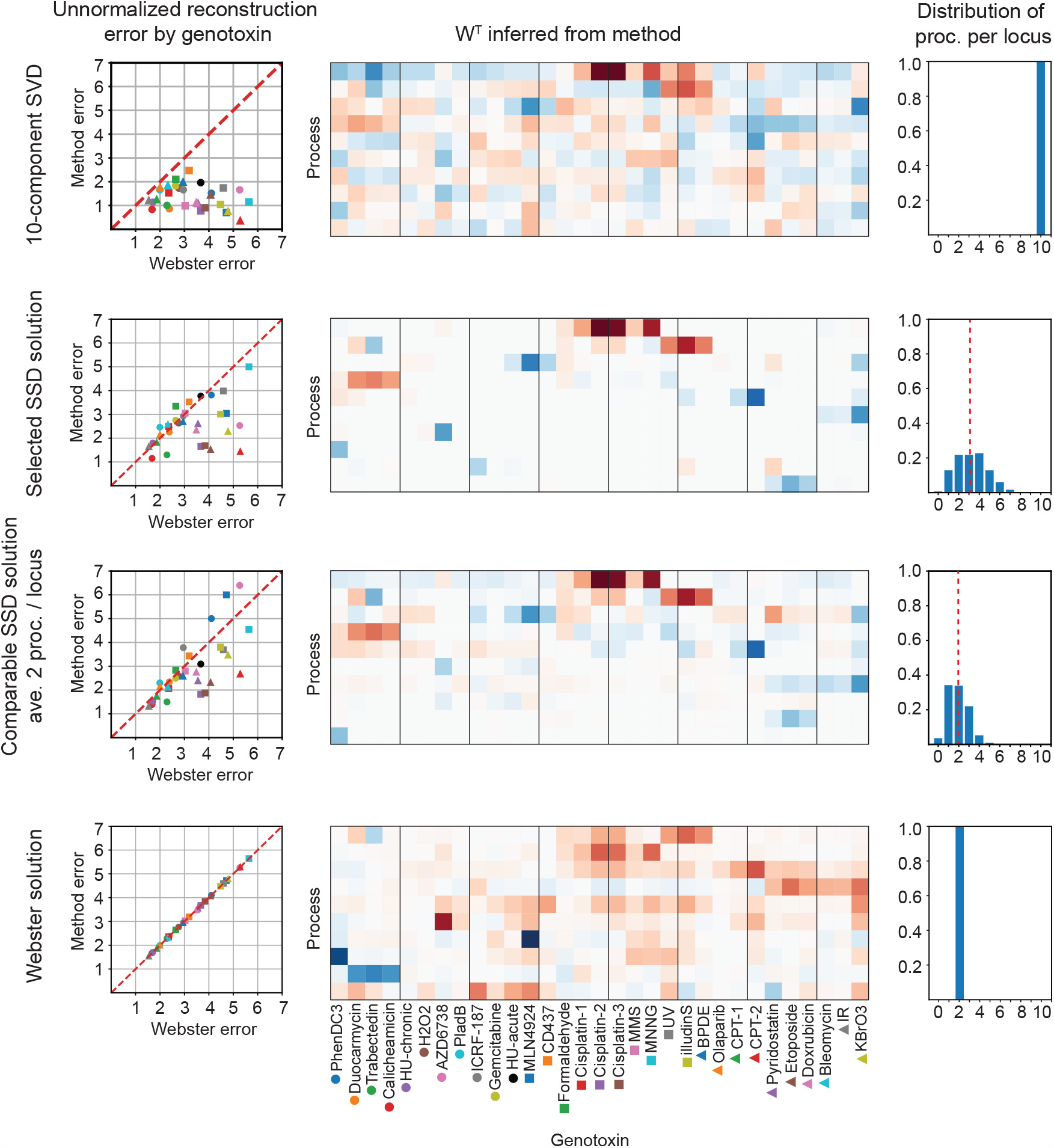
Comparison of SVD, SSD, and Webster decompositions on genotoxin data. Each row corresponds to a decomposition found by SVD, SSD, or Webster, as labeled on the left. The leftmost column compares the unnormalized reconstruction error for each method as compared to Webster for each genotoxin separately. The center column illustrates the process-genotoxin map found by each method; the SSD solutions exhibit the most sparsity. The rightmost column illustrates the distribution of processes per locus in each decomposition. The red dotted line depicts the mean.

**FIG. S7:**
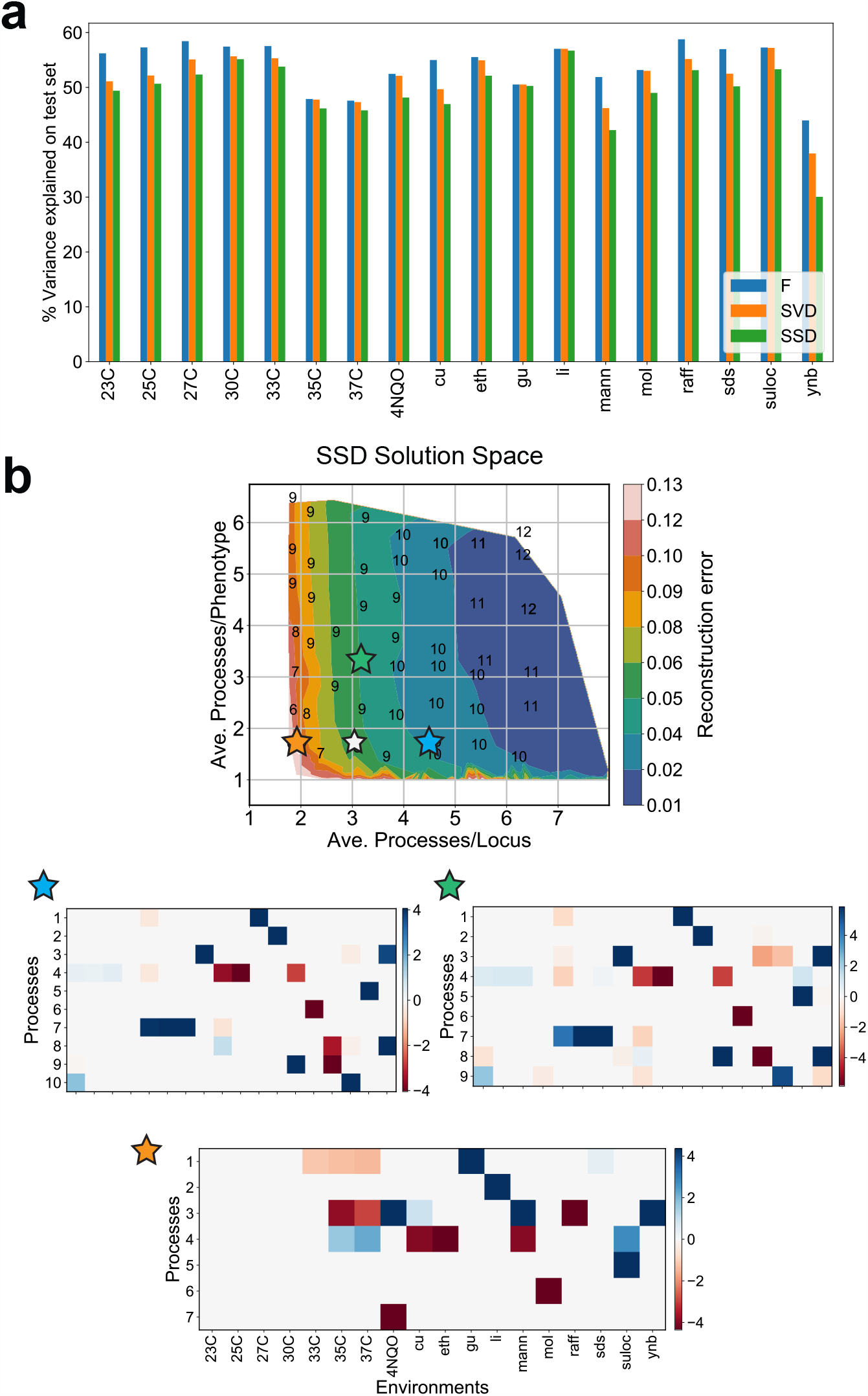
Variance explained on the test set and the process-phenotype map of other SSD solutions for the yeast cross data. (a) The percentage variance explained when predicting the fitness in individual environments on a test set of genotypes (i.e., as 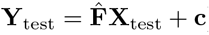), shown here when 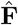 is the full additive effects matrix **F** (blue), the 8-component SVD approximation of **F** (orange) and the 8-component SSD solution analyzed in the main text and marked in Figure 5d (green). (b) The process-phenotype map **W** for three additional solutions marked in the solution space. The white star marks the SSD solution discussed in the main text. Note that the general features are conserved between the solutions marked with the blue and green stars. The much sparser solution marked by the orange star also devotes dedicated processes for li, gu and mol, but tends to group the other processes together.

